# Inhibition of an immunometabolic axis of mTORC1 activation extends mammalian healthspan

**DOI:** 10.1101/2023.07.09.548250

**Authors:** Anissa A. Widjaja, Wei-Wen Lim, Sivakumar Viswanathan, Sonia Chothani, Ben Corden, Joyce Wei Ting Goh, Jessie Tan, Chee Jian Pua, Radiance Lim, Brijesh K. Singh, Dasan Mary Cibi, Susanne Weber, Sze Yun Lim, Eleonora Adami, Benjamin L. George, Mark Sweeney, Chen Xie, Madhulika Tripathi, Dominic J Withers, Norbert Hübner, Sebastian Schafer, Lena Ho, Jesus Gil, David Carling, Stuart A. Cook

**Affiliations:** Cardiovascular and Metabolic Disorders Program, Duke-National University of Singapore Medical School, Singapore; National Heart Research Institute Singapore, National Heart Centre Singapore, Singapore; Barts Heart Centre, Barts Health NHS Trust, London, UK; VVB Bio Pte Ltd, Singapore; Cardiovascular and Metabolic Sciences, Max Delbrück Center for Molecular Medicine in the Helmholtz Association (MDC), Berlin, Germany; MRC-London Institute of Medical Sciences, Hammersmith Hospital Campus, London, UK; Institute of Clinical Sciences, Faculty of Medicine, Imperial College, London, UK; DZHK (German Centre for Cardiovascular Research), Partner Site Berlin, Berlin, Germany; Charité-Universitätsmedizin, Berlin, Germany

## Abstract

Human ageing is associated with metabolic dysfunction, sarcopenia and frailty that taken together reduce healthspan. For age-associated diseases and lifespan, ERK, AMPK and mTORC1 represent critical pathways, across species^1–7^. Here we examined the hypothesis that IL11, recently shown to regulate ERK/mTORC1, is an inflammaging factor important for healthspan. As mice age, IL11 is progressively upregulated in liver, skeletal muscle, and fat to stimulate an ERK/AMPK/mTORC1 axis of cellular, tissue- and organismal-level ageing pathologies. In old mice, deletion of *Il11* or *Il11ra1* protects against metabolic multi-morbidity, sarcopenia, and frailty. Administration of anti-IL11 therapy to elderly mice for six months reactivates an age-repressed program of white fat beiging, reverses metabolic dysfunction, restores muscle function, and reduces frailty. Across studies, inhibition of IL11 lowers epigenetic age, reduces telomere attrition, and preserves mitochondrial function. Towards clinical translation, we generated, humanised, and engineered a neutralising, high-affinity IL11 antibody. These studies identify IL11 as a key inflammaging factor and therapeutic target for mammalian healthspan.

## Introduction

The major evolutionarily conserved signalling mechanisms that regulate lifespan across species include ERK, STK11 (LKB1), AMPK, mTORC1, and IGF1/insulin modules^1–4^. These pathways are perturbed by dysregulated nutrient sensing to cause mitochondrial dysfunction and cellular senescence, comprising driver hallmarks of ageing^1^. In old age, the AMPK/mTORC1 axis is uniquely important for metabolic health, with notable effects in adipose tissue^8–10^. Chronic systemic inflammation “inflammaging” is another important hallmark of ageing, predicts mortality in the elderly and, along with fibrosis, is universally associated with senescence^1, 11^.

Ageing studies to date have focused mostly on lifespan extension that can be achieved at the expense of healthspan, merely represent extended periods of frailty and, in mice, primarily reflects reduced murine cancers^12–14^. For these reasons, there is a need for studies beyond lifespan: to better understand healthspan. Laboratory mice are particularly suited for healthspan experiments as ageing pathologies important for human wellbeing and function, such as frailty and sarcopenia, are readily apparent and translatable^10, 13, 15–17^.

IL11 has no known function in ageing but is related to IL6, an inflammageing factor^18^, and was recently shown to be pro-inflammatory and pro-fibrotic and to activate ERK, inhibit LKB1/AMPK and stimulate mTORC1 across mouse and human cell types (**Fig. 1a**)^19–21^. Conditional deletion of *Il11ra1* in epithelial cells is permissive for tissue regeneration across organs, which is impaired in the elderly^22, 23^. As an immunometabolic factor, IL11 causes liver steatosis, visceral adiposity and impaired muscle cell function, known features of ageing^24, 25^. IL11 is also a component of the senescence associated secretory phenotype (SASP) and can cause senescence^6, 26, 27^. Here, using genetic and pharmacologic approaches, we explore the hypothesis that IL11 is an inflammaging factor and a therapeutic target for preventing and treating multi-morbidity in old age to extend healthspan.

**Figure 1.**
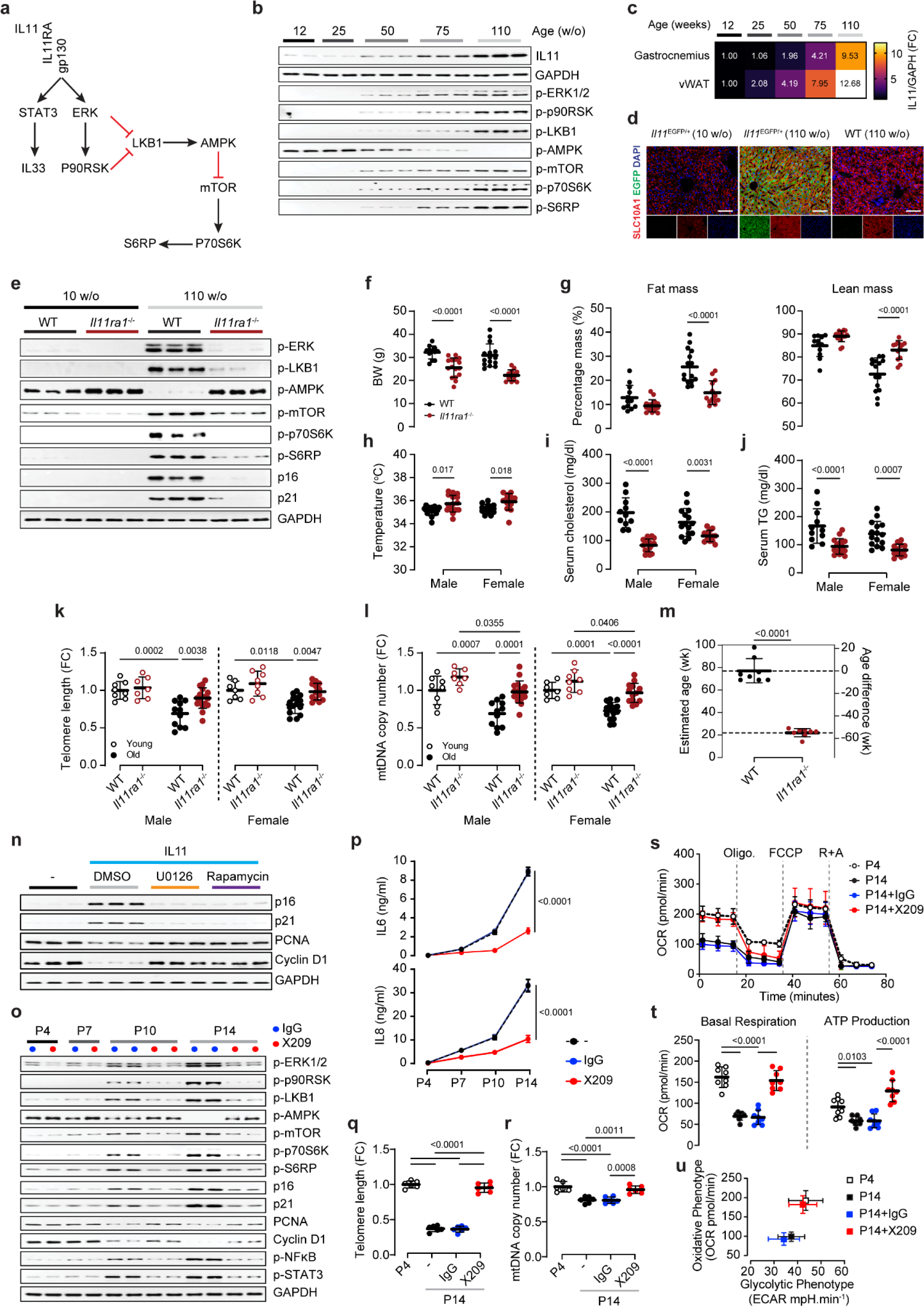
The IL11/ERK/mTORC1 signalling module is upregulated in ageing to cause senescence and metabolic decline. **a** Schematic showing signalling pathways by which IL11 induces LKB/AMPK inactivation and mTOR activation. **b** Western blots (WB) of IL11, GAPDH, p-ERK1/2, p-p90RSK, p-LKB1, p-AMPK, p-mTOR, p-p70S6K, p-S6RP in livers from 12, 25, 50, 75, and 110-week-old (w/o) male mice (n=5/group). **c** Heat map showing densitometry of IL11 protein expression normalised to GAPDH in gastrocnemius and visceral white adipose tissues (vWAT) from 12 to 110 w/o male mice (n=5/group). **d** Representative immunofluorescence images (scale bars, 100 µm) of EGFP and SLC10A1 expression in the livers of 10 and 110 w/o *Il11*-*EGFP* mice (representative dataset from n=3/group). **e** WB of p-ERK1/2, p-LKB1, p-AMPK, p-mTOR, p-p70S6K, p-S6RP, p16, p21, and GAPDH in livers from 10 and 110 w/o male wild-type (WT) and *Il11ra1^-/-^* mice (n=3/group). **f** Body weights (BW), **g** percentages of fat and lean mass (normalised to BW), **h** body temperatures, and the levels of **i** serum cholesterol, and **j** serum triglycerides (TG) of 110 w/o male and female WT and *Il11ra1^-/-^* mice (male WT for (**f-h**), n=12; male WT for (**i-j**), n=11; male *Il11ra1^-/-^*, n=16; female WT, n=15; female *Il11ra1^-/-^*, n=13). Liver **k** telomere length and **l** mitochondria DNA (mtDNA) copy number from young (10 w/o) and old (110 w/o) male and female WT and *Il11ra1^-/-^* mice (young male WT, n=8; young male *Il11ra1^-/-^*, n=7; old male WT, n=11; old male *Il11ra1^-/-^*, n=17; young female WT, n=7; young female *Il11ra1^-/-^*, n=8; old female WT, n=15; old female *Il11ra1^-/-^*, n=13). **m** Estimated liver DNA methylation age from male 110 w/o WT and *Il11ra1^-/-^* mice (n=8/group). **n** Effects of U0126 (10 µM) and rapamycin (10 nM) on p16, p21, Cyclin D1, and PCNA protein expression in IL11 (5 ng/ml)-stimulated HCFs by WB (n=6/group). **o-u** Data for HCF passage 4 (P4), 7, 10, and 14 that had been passaged in the presence of either IgG or anti-IL11RA (X209; 2µg/ml) from P2. **o** WB of p-ERK1/2, p-p90RSK, p-LKB1, p-AMPK, p-mTOR, p-p70S6K, p-S6RP, p-NFκB, p-STAT3, p16, p21, PCNA, Cyclin D, and GAPDH, **p** IL6 and IL8 levels in the supernatant based on ELISA, **q** telomere length, and **r** mtDNA copy number (n=6/group). Seahorse assay showing **s** mitochondrial oxygen consumption rate (OCR), **t** changes in OCR during basal respiration and ATP production states, and **u** oxidative and glycolytic energy phenotypes at baseline (n=8/group). **f-m, p-u** Data are shown as meanLJ±LJSD. **f-l** Two-way ANOVA with Sidak’s correction; **m** two-tailed Student’s t-test; **p** two-way ANOVA, **q, r, t** one-way ANOVA with Tukey’s correction. FC: fold change.

## Results

### IL11 is upregulated in metabolic tissues with age

We determined IL11 expression in the liver, visceral gonadal white adipose tissue (referred to as vWAT from here onwards) and skeletal muscle (gastrocnemius) in mice over a time course of ageing, which revealed progressive upregulation of IL11 in all these tissues (**Fig. 1b, c; Extended Data Fig. 1a**). IL11 upregulation was confirmed in livers, vWAT and skeletal muscle of 2-year-old male and female mice (**Extended Data Fig. 1b**). With age, there was progressive activation of ERK/p90RSK, inactivation of LKB1/AMPK and mTOR/p70S6K activation, which comprise the IL11 signalling module, in liver and muscle (**Fig. 1a, b; Extended Data Fig. 1c, d**).

To identify the cell types expressing *Il11* in old mice we queried the Tabula Muris Senis^28^, which was uninformative as *Il11* is lowly expressed and IL11 levels are translationally regulated^21^. We thus aged *Il11:EGFP* reporter mice^22^ and performed immunohistochemistry. In two-year-old *Il11:EGFP* mice, IL11 was most apparent in parenchymal cells (hepatocytes, liver; adipocytes, vWAT; myocytes, skeletal muscle), with additional expression seen in stromal, epithelial cells and nerves across tissues (**Fig. 1d; Extended Data Fig. 1e-g**).

### IL11 activates ERK/mTOR and increases senescence markers with age

To explore the functional relevance of IL11 upregulation we studied 10-week-old (w/o) and two-year-old mice deleted for *Il11ra1* (*Il11ra1^-/-^*) and wildtype littermate controls (WT). On immunoblots of liver extracts, as compared to young mice, old WT mice had increased phospho (p)-ERK, p-p90RSK, diminished LKB1/AMPK activity and increased levels of p-mTOR, p-p70S6K and p-S6RP (**Fig. 1e; Extended Data Fig. 2a**). Levels of the canonical senescence markers p16(Inka4) and p21(Waf1/Cip1) were increased in livers of old WT mice, as expected. In contrast, kinase phosphorylation levels in livers of old *Il11ra1^-/-^*mice were similar to those of young mice and there was no increase in p16 or p21. These signalling and senescence marker data were replicated in skeletal muscle and vWAT from *Il11ra1^-/-^*male mice (**Extended Data Fig. 2b, c**).

### IL11 causes metabolic decline in old mice

As compared to WT, two-year-old *Il11ra1^-/-^* mice of both sexes had lower body weights (**Fig. 1f**) and female *Il11ra1^-/-^*mice had decreased fat mass and increased lean mass by body composition analysis (**Fig. 1g**). Old *Il11ra1^-/-^* mice of both sexes had slightly higher body temperatures than WTs (**Fig. 1h**).

In *ex vivo* studies, old female *Il11ra1^-/-^* mice had lower indexed vWAT mass and both sexes of *Il11ra1^-/-^* mice had elevated indexed gastrocnemius mass (**Extended Data Fig. 2d**). Liver indices were similar between genotypes whereas liver triglyceride levels were lower in *Il11ra1^-/-^*mice (**Extended Data Fig. 2d, e**). In both sexes of old WT mice, serum cholesterol and triglyceride levels were higher, whereas serum levels of β-hydroxybutyrate (BHB), which is advocated as an anti-ageing supplement^29^, were lower than those of *Il11ra1^-/-^* littermates (**Fig. 1i, j**; **Extended Data Fig. 2f**)

Livers of old *Il11ra1^-/-^* mice of both sexes had reduced age-dependent expression of pro-inflammatory (*Ccl2*, *Ccl5*, *Tnf*α, *Il1b* and *Il6*) and fatty acid synthesis (*Acc*, *Fasn* and *Srebp1c)* genes (**Extended Data Fig. 2g, h**). Furthermore, serum alanine aminotransferase (ALT) and aspartate aminotransferase (AST) levels, markers of hepatocyte damage, were raised in old WT mice but not in old *Il11ra1^-/-^*mice (**Extended Data Fig. 2i, j**).

### Ageing biomarkers are associated with IL11 activity

Biomarkers most strongly associated with biological age include epigenetic clocks, telomere length and mitochondrial DNA (mtDNA) copy number^1, 30^. We assessed telomere lengths and mtDNA copy numbers in the livers and skeletal muscle of both young and two-year-old mice of both sexes and found both phenotypes to be preserved in livers of old *Il11ra1^-/-^* mice, which was replicated in skeletal muscle (**Fig. 1k, l; Extended Data Fig. 2k, l**). We then used a liver-specific epigenetic clock to predict the ages of two-year-old *Il11ra1^-/-^*and WT mice: those of *Il11ra1^-/-^* genotype were estimated to be 55 weeks younger than WT (**Fig. 1m**).

### The IL11/ERK/mTORC1 signalling module causes senescence

Senescence is widely recognised as a key driver of ageing pathologies across cells, tissues and species and IL11 was recently shown to induce senescence in stromal and epithelial cells^6, 11, 26, 27^. We thus explored the role of IL11 in senescence in greater detail in primary human cells.

Stimulation of fibroblasts with IL11 activated the ERK/mTOR signalling module, increased p16/p21 levels and reduced PCNA/Cyclin D1 expression (**Fig. 1n; Extended Data Fig. 3a**). IL11-induced fibroblast senescence was prevented by MEK/ERK (U0126) or mTORC1 (rapamycin) inhibitors (**Fig. 1n**), consistent with the role of these pathways in senescence^6, 27^. Supernatants of IL11 stimulated fibroblasts had increased levels of SASP factors (IL6 and IL8) that were inhibited by U0126 or rapamycin (**Extended Data Fig. 3b, c**).

IL11 expression was increased in hepatocytes with age (**Fig. 1d**) but it was unknown if IL11 causes hepatocyte senescence. We profiled mTORC1-dependent SASP factors (IL6, IL8, LIF, VEGFA, HGF, CCL2, CXCL1, CXCL5, CXCL6 and CCL20^6^) in supernatants of hepatocytes stimulated with IL11 for 6 or 24 hours and observed significant upregulation of the majority of these proteins (**Extended Data Fig. 3d**). As seen in fibroblasts, IL11 caused activation of the IL11 signalling module in hepatocytes and this was associated with ERK- and mTORC1-dependent senescence gene upregulation and proliferation gene downregulation (**Extended Data Fig. 3e**). IL11-induced IL6 and IL8 secretion from hepatocytes and ERK- and mTORC1-dependence of this was shown (**Extended Data Fig. 3f, g**).

### Replicative senescence is IL11-dependent

Replicative senescence (RS) is thought important in ageing and to understand if IL11 regulates this pathobiology, we modelled RS in primary human fibroblasts serially passaged (from P4 to P14) in the presence or absence of a neutralising IL11RA antibody (X209)^22^. There was passage-dependent activation of the IL11 signalling module, which was associated with increased p16/p21, diminished PCNA/Cyclin D1, and elevated pro-inflammatory markers (p-NFκB and p-STAT3) (**Fig. 1o**).

In the presence of X209, the IL11 signalling module, features of senescence, cell cycle withdrawal, and inflammation were collectively reduced (**Fig. 1o; Extended Data Fig. 3h**). Levels of IL8, IL6 and IL11 steadily increased in supernatants of later passages and this was inhibited by X209 (**Fig. 1p; Extended Data Fig. 3i**). Paracrine amplification of senescence occurs via an mTORC1-dependent SASP process^6^ and we tested whether IL11 contributed to this phenomenon. Incubation of early passage (P4) fibroblasts with medium from late passage (P14) fibroblasts resulted in IL11-dependent upregulation of p16 and p21 in the P4 cells, indicating a role for IL11 in paracrine senescence (**Extended Data Fig. 3j**).

### Inhibition of IL11 reduces ageing biomarkers and improves mitochondrial function

Ageing biomarkers are apparent, in part, in cells undergoing RS^1, 31^. We profiled fibroblasts at P4 and at P14 and found that telomere lengths and mtDNA copy numbers were similar between P4 and P14+X209 cells, whereas P14+IgG cells had reduced telomere lengths and mtDNA copy numbers (**Fig. 1q, r**). In keeping with this, basal metabolic respiration, ATP production and the basal energy phenotype were impaired in P14+IgG fibroblasts whereas P14+X209 cells were indistinguishable from P4 fibroblasts (**Fig. 1s-u**).

### *Il11*^-/-^ mice are protected from age-associated obesity and frailty

Our initial data in *Il11ra1*-deleted mice and human cells pointed to an important role for IL11 in ageing, which we dissected in young (3-month-old) and aged (2-year-old) female mice deleted for *Il11 (Il11*^-/-^) and WT littermates^32^. This allowed for data replication across genotypes and strains with loss-of-function in IL11 signalling^32^.

Immunoblots on protein extracts from liver, fat and skeletal muscle of young and old *Il11*^-/-^ and WT mice confirmed IL11 upregulation in this additional strain in old age, while also verifying the specificity of the anti-IL11 antibody (X203) (**Fig. 2a**). Old female *Il11*^-/-^ mice appeared leaner, had reduced body weights (27% lower) and fat mass (72% lower) along with preserved lean mass, as compared to old WT mice (**Fig. 2b-d**).

**Figure 2.**
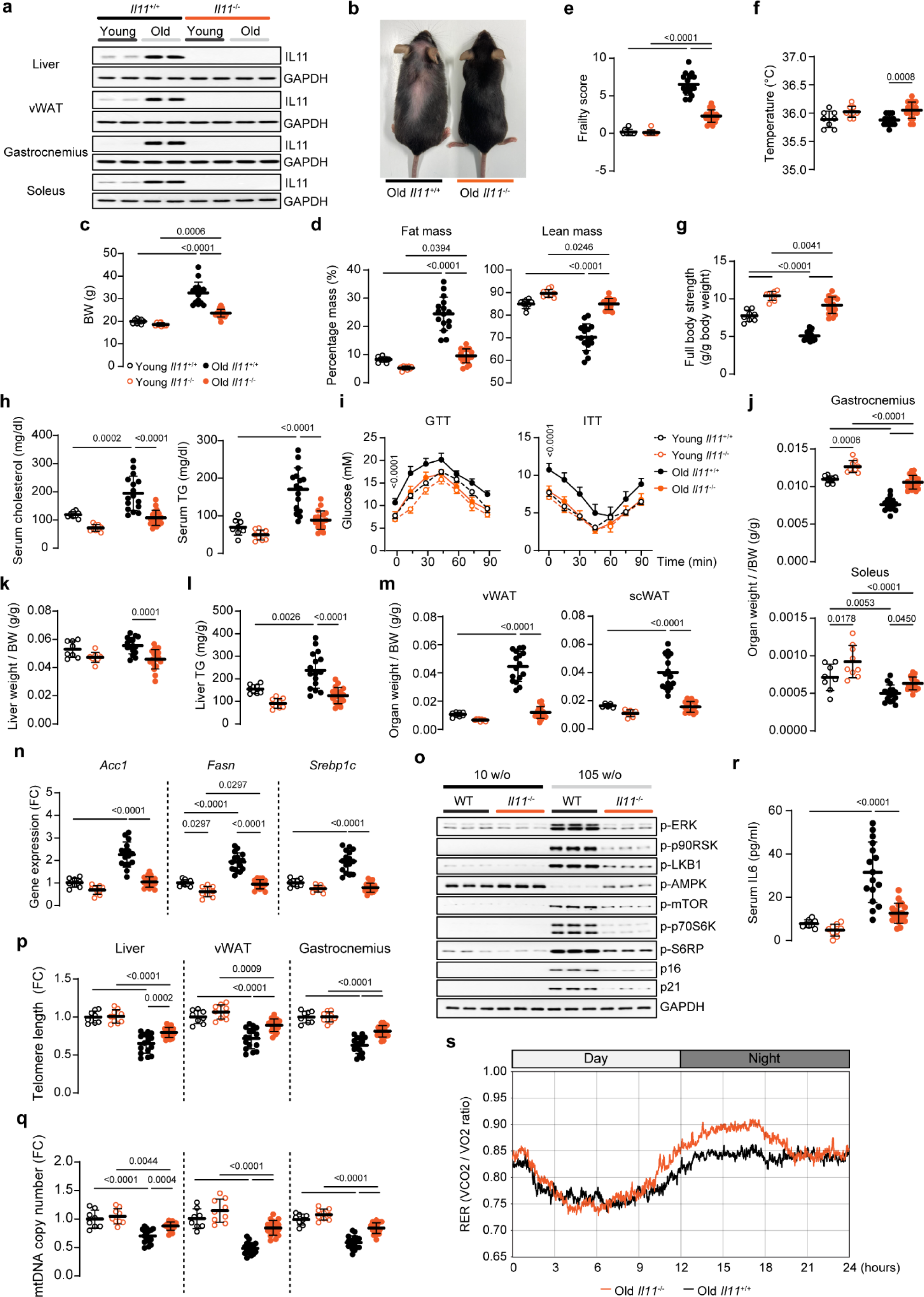
Female *Il11*-deleted mice are protected from age-associated obesity, frailty, and metabolic decline. **a** WB of IL11 and GAPDH expression from the indicated organs of young (12 w/o) and old (105 w/o) male WT and *Il11^-/-^* mice (n=2/group). **b** Representative picture of 108 w/o female WT and *Il11^-/-^* mice. **c** Body weights (BW), **d** percentages of fat and lean mass (normalised to BW), **e** frailty scores, **f** body temperatures, **g** full body grip strength measurements, **h** serum cholesterol and TG levels, **i** glucose and insulin tolerance tests (GTT and ITT) from young (12 w/o) and old (105 w/o) female WT and *Il11^-/-^* mice. Weights of **j** skeletal muscle (gastrocnemius and soleus), and **k** liver (indexed to BW), **l** liver TG levels, **m** indexed weights of vWAT and inguinal subcutaneous WAT (scWAT), **n** relative vWAT mRNA expression levels of *Acc1*, *Fasn*, and *Srebp1c*, **o** WB showing activation status of ERK1/2, p90RSK, LKB1, AMPK, mTOR, p70S6K, S6RP and protein expression levels of p16, p21, and GAPDH (representative datasets from n=6/group) in vWAT, **p** telomere length and **q** mtDNA copy number, **r** serum IL6 levels from young and old female WT and *Il11^-/-^* mice. **s** Respiratory exchange ratio (RER) measurements in 68-70 w/o male WT and *Il11^-/-^* mice (n=10/group). **c-n, p-r** Data are shown as meanLJ±LJSD (young WT, n=8; young *Il11^-/-^*, n=9; old WT, n=16; old *Il11^-/-^*, n=18). **c-h, j-n, p-r** Two-way ANOVA with Sidak’s correction; **i** two-way ANOVA. FC: fold change.

The mean frailty score, as measured using a 27-point score^33^, of old female *Il11*^-/-^ mice was noticeably lower than that of WT mice and body temperatures were mildly elevated (**Fig. 2e, f**). The lower frailty scores were driven by improvements across a range of phenotypes including tremor, loss of fur colour, gait disorders and vestibular disturbance (**Extended Data Table. 1**). Muscle strength, an important measure of sarcopenia and frailty in humans^15, 18^, was higher in both young and old *Il11^-/-^* mice, as compared to age-matched controls (**Fig. 2g; Extended Data Fig. 4a**).

### Deletion of *Il11* improves metabolism in old age

Chronic inhibition of mTOR with rapamycin, although targeted to raptor/mTORC1, can cause glucose intolerance due to indirect inhibition of mTORC2^34, 35^. It was therefore important to more fully assess the effects of Il11 inhibition on metabolism and glucose utilisation in old mice, to exclude side effects.

As mice aged, there were increases in serum ALT, AST, cholesterol, and triglycerides and decreases in BHB in WT mice that were collectively reduced in old *Il11*^-/-^ mice (**Fig. 2h; Extended Data Fig. 4b, c**). Fasting blood glucose (FBG) concentrations were lower in old *Il11^-/-^* mice as compared to WT controls (**Fig. 2i**). Glucose and insulin tolerance tests (GTT and ITT) profiles and areas under the curve (AUC) of old *Il11^-/-^* mice were similar to young WT mice, whereas GTTs and ITTs of old WT mice were impaired (**Fig. 2i; Extended Data 4d**). Even in young adulthood, GTTs of *Il11^-/-^* mice were better than in WT. Hence, chronic inhibition of Il11 has metabolic benefits, unlike prolonged use of rapamycin.

### Reduced adiposity, mTORC1 activity, and ageing biomarkers in old *Il11^-/-^*mice

In human GWAS, nsSNPs in *IL11* are associated with a small reduction in height^36^. We measured nose-rump lengths and found old *Il11^-/-^* mice to be ∼5% shorter than WT (**Extended Data Fig. 4e**). Age-dependent sarcopenia was apparent in both old female WT and *Il11^-/-^* mice but with overall greater indexed muscle weights seen in *Il11^-/-^* mice (**Fig. 2j**).

As compared to old WT, old *Il11^-/-^* mice had reduced liver mass indices and liver triglyceride content (**Fig. 2k, l**). Furthermore, indexed vWAT and inguinal subcutaneous WAT (scWAT) masses were diminished (73% and 61%, respectively) in old *Il11^-/-^* mice, whereas brown adipose tissue (BAT) mass was unchanged (**Fig. 2m; Extended Data Fig. 4f**)^8, 10^. Increased lipolysis was excluded as an underlying cause of reduced WAT given IL11 inhibits this^37^. We profiled expression of fatty acid synthesis genes in vWAT and found their expression to be increased with age in old WT mice but not in old *Il11^-/-^* mice (**Fig. 2n**).

We assessed the IL11/mTORC1 signalling axis in vWAT and gastrocnemius of old WT mice and replicated its activation and association with senescence, which were not the case in old *lI11^-/-^* mice (**Fig. 2o; Extended Data Fig. 4g, h**). Pro-inflammatory gene expression (*Ccl2, Ccl5, Tnf*ll!*, Il1*L and *Il6*) was increased in vWAT of old WT mice, as compared to young mice, but this was diminished in vWAT of old *lI11^-/-^* mice (**Extended Data Fig. 4i**).

As compared to young mice, telomere lengths and mtDNA content of liver, skeletal muscle and vWAT were reduced in old WT mice and these effects were attenuated in old *lI11^-/-^* mice (**Fig. 2p, q**). Serum IL6 levels, a biomarker that predicts frailty and mortality in the elderly^18^, were largely elevated in old WT mice but not in old *Il11^-/-^* mice (**Fig. 2r**).

### Male IL11 knockout mice are protected from age-associated disease

For completeness, we repeated in life and organ-level phenotypic experiments in young and old male WT and *Il11^-/-^* mice. Age-related changes in body habitus, body weight, fat mass and lean mass were mitigated in male *Il11*^-/-^ mice (**Extended Data Fig. 5a-c**). Frailty scores and body temperatures of old male *Il11*^-/-^ mice were lower and higher, respectively than those of WT male mice (**Extended Data Fig. 5d, e and Extended Data Table 2**). Lower frailty scores were driven by improvements in coat colour and condition, tremor, vestibular disturbance and vision loss. Indexed full body and forepaw grip strengths were higher in *Il11^-/-^* mice as compared to WT (**Extended Data Fig. 5f**). In old *Il11^-/-^* mice, FBG levels were lower than old WT and dynamic profiles and AUC of GTTs and ITTs were similar to young mice (**Extended Data Fig. 5g**). Noticeably, GTTs and ITTs in young *Il11^-/-^* mice were improved as compared to young WT mice.

To characterise whole body metabolism more fully, we housed 80 w/o male mice in metabolic cages. The respiratory exchange ratio (RER) was higher in *Il11^-/-^* mice as compared to WT mice (**Fig. 2s; Extended Data Fig. 5h**) with aged mice of both genotypes having expectedly lower RERs than young mice during the dark phase^17, 38^. After a period of starvation, refeeding resulted in a greater increase of RER in *Il11^-/-^* mice compared to WT mice, consistent with better metabolic flexibility (**Extended Data Fig. 5h**)^39^. While *Il11^-/-^* mice were leaner and weighed less than WT, they consumed more food and had similar levels of activity (**Extended Data Fig. 5h**).

Body lengths of old male *Il11^-/-^* mice were slightly shorter than old male WT (**Extended Data Fig. 5i**). Age-dependent sarcopenia was apparent in old WT mice but less so in old *Il11^-/-^* mice and indexed liver weights were similar between genotypes (**Extended Data Fig. 5j, k**). An incidental finding of enlarged seminal vesicles, a recognised ageing phenomenon^40^, was more common in old WT as compared to old *Il11^-/-^*mice (WT: 11/15 and *Il11^-/^*: 1/14; P=0.0003).

Once again, the most notable organ difference associated with IL11 loss-of-function in old mice was seen in WAT, with a 73% and 43% reduction in the indexed mass of vWAT and scWAT, respectively in old *Il11^-/-^* mice as compared to WT (**Extended Data Fig. 5l**). Indexed BAT mass was similar across genotypes and ages, as seen in females.

### Therapeutic inhibition of IL11 has pleiotropic benefits in old mice

Lifelong lack of IL11 signalling in mice deleted for *Il11ra1* or *Il11* may confer developmental effects and was shown to impact some aspects of metabolism in young mice (**Fig. 1**, **2**). To exclude early life events as a cause of late life phenotypes and to explore the translational relevance of our findings, we administered a neutralising IL11 antibody (X203) or IgG control to aged mice (**Fig. 3a**)^22^.

**Figure 3.**
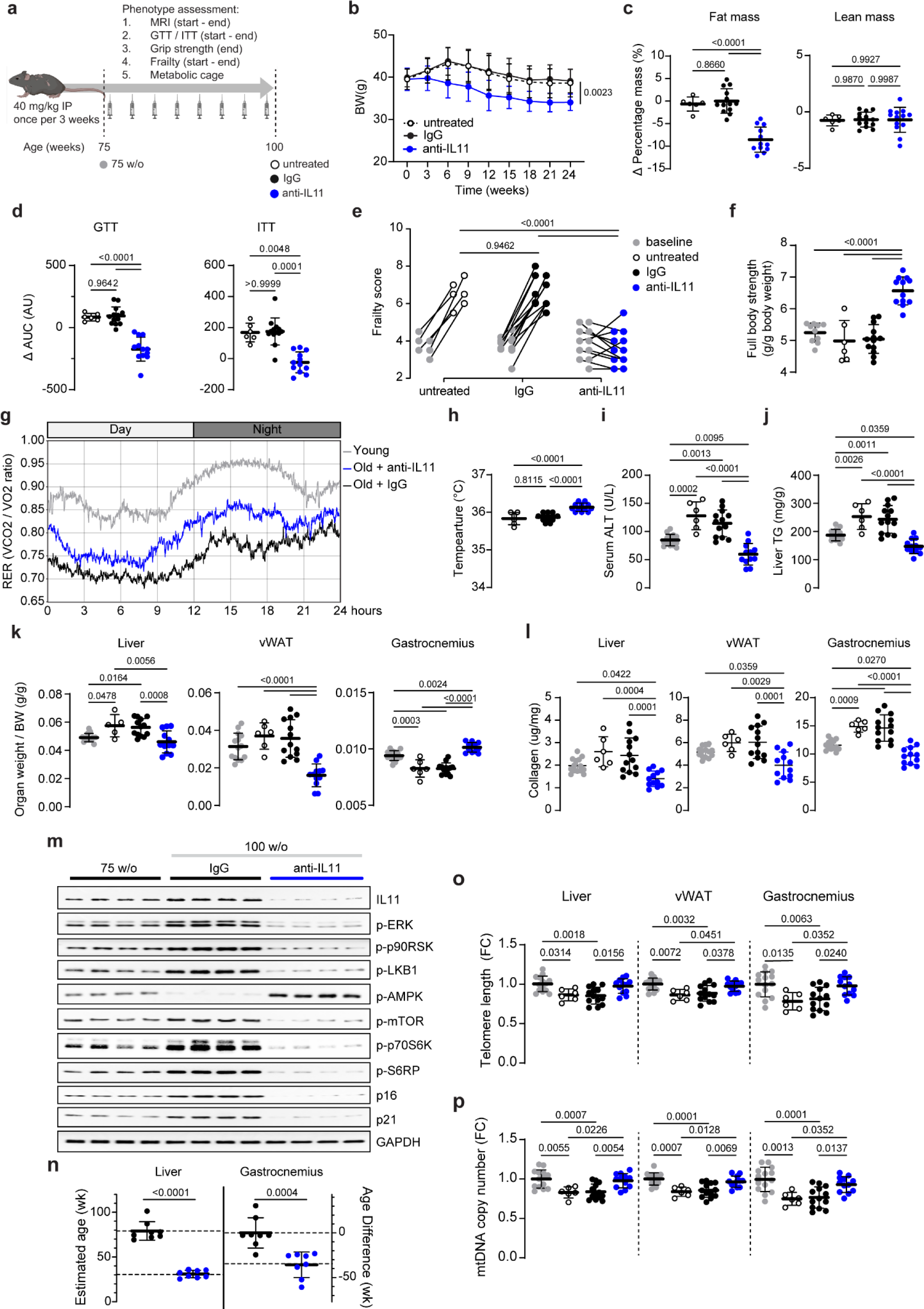
Therapeutic inhibition of IL11 reverses age-associated metabolic dysfunction, pathogenic signalling and sarcopenia. **a** Schematic of anti-IL11 (X203) therapeutic dosing experiment in old male mice for experiments shown in (**b-p**). Mice were either aged naturally (untreated) or given either X203 or an IgG control antibody (40 mg/kg, every 3 weeks) starting from 75 weeks of age for a duration of 25 weeks. **b** Body weights across time. **c-f** Changes (Δ; values at end-point (100 w/o) - values at starting point (75 w/o)) in **c** fat and lean mass percentage, and **d** area under the curve (AUC) of GTT and ITT. **e** Frailty scores at starting and end-point. **f** Full body grip strength. **g** RER measurements in young (14 w/o) and IgG/X203-treated old (81 w/o) mice - 6 weeks after IgG/X203 administration was started (n=10/group). **h** Body temperatures, **i** serum ALT level, **j** liver TG levels. **k** indexed weights of and **l** total collagen content (by hydroxyproline assay) in liver, gastrocnemius, and vWAT. **m** WB showing activation status of ERK1/2, p90RSK, LKB1, AMPK, mTOR, p70S6K, S6RP and protein expression levels of IL11, p16, p21, and GAPDH in vWAT (representative datasets from n=6/group). **n** Estimated liver and gastrocnemius DNA methylation age, **o** telomere length and **p** mtDNA copy number. **b-d, f, h-l, n-p** Data are shown as meanLJ±LJSD, **e** data are shown as values recorded at starting and end-point (75 w/o control, n=14; untreated 100 w/o, n=6; IgG-treated 100 w/o n=13; X203-treated 100 w/o, n=12). **b** Two-way ANOVA, **c-f, h-l, o-p** one-way ANOVA with Tukey’s correction, **n** two-tailed Student’s t-test. FC: fold change; AU: arbitrary units.

As compared to IgG controls, mice receiving X203 from 75-to-100 w/o progressively lost body weight that was defined by a reduction in indexed fat mass (**Fig. 3b, c**). Impaired glucose metabolism was apparent across experimental groups at study start and was reversed in 100 w/o mice receiving X203, whereas IgG had no effect (**Fig. 3d**).

Frailty scores were mildly elevated across aged experimental groups at study initiation (**Fig. 3e**). Over the 25-week study period, mice receiving no treatment or IgG exhibited frailty progression, whereas those receiving X203 showed no increase (**Fig. 3e**; **Extended Data Table 3**). Muscle strengths of 100 w/o mice receiving anti-IL11 were higher as compared to those receiving IgG and also when compared to 75 w/o mice, showing reversal of muscle dysfunction (**Fig. 3f; Extended Data Fig. 6a**).

After 6 weeks of antibody administration, mice were studied in metabolic cages. As compared to those receiving IgG, the RER of the X203 group was higher overall but lower when directly compared to a cohort of young mice (**Fig. 3g; Extended Data Fig. 6b**), suggesting that X203 slows the trend of age-associated RER decline. Administration of X203 was associated with increased food intake whilst activity levels were similar between study groups (**Extended Data Fig. 6b**). Core temperatures of X203-treated mice were higher than age-matched untreated mice and those mice receiving IgG (**Fig. 3h**).

Mice left untreated or given IgG had elevated serum cholesterol, triglycerides and IL6 that were collectively lowered, below 75 w/o levels, by X203 in 100 w/o mice (**Extended Data Fig. 6c**). The reciprocal was true for serum BHB levels (**Extended Data Fig. 6d**). Over the course of the experiment, markers of liver damage, hepatic triglyceride content, and indexed liver mass increased in untreated and IgG control mice, whereas these phenotypes were either reversed or reduced in mice receiving X203 (**Fig. 3i-k; Extended Data Fig. 6e**).

There was a notable reduction in indexed vWAT mass and an increase in indexed muscle mass in 100 w/o mice receiving X203, as compared to both age-matched controls and 75 w/o mice, indicating a reversal of age-related visceral adiposity and sarcopenia (**Fig. 3k; Extended Data Fig. 6f**). Mice receiving X203 had diminished scWAT, as compared untreated age matched controls and those administered IgG (**Extended Data Fig. 6g**). In contrast to *Il11^-/-^* mice, administration of X203 was associated with a small increase in indexed BAT mass (**Extended Data Fig. 6g**). The ageing phenotype of enlarged seminal vesicles was again more commonly seen in mice with IL11 loss-of-function (IgG: 8/13 and X203: 2/12; P=0.022)^40^.

### Anti-IL11 reverses age-associated fibrosis, signalling and biomarkers

Fibrosis is a canonical feature of ageing and a hallmark of senescence and IL11 is known to be pro-fibrotic in human cells and in mouse models of human disease^1, 11, 41^. We quantified fibrosis biochemically in vWAT, skeletal muscle, and livers of old mice across experimental groups. There was reversal of tissue fibrosis in all organs of mice receiving X203, as compared to 75 w/o mice, that was not seen in 100 w/o untreated or IgG control groups (**Fig. 3l**).

We profiled the IL11 signalling in vWAT. As compared to 75 w/o mice, those receiving IgG had further activation of the IL11/mTORC1 axis and higher expression of senescence markers (**Fig. 3m; Extended Data Fig. 6h**). In contrast, mice receiving X203 had reduced IL11 signalling activity, as compared to either IgG controls or 75 w/o mice (**Fig. 3m; Extended Data Fig. 6h**). This shows reversal of the canonical age-associated signalling pathologies (ERK and mTOR activation, AMPK inactivation) with the administration of anti-IL11 to old mice.

Epigenetic clock studies showed ∼30 and ∼50-week reductions in clock-derived age estimates in skeletal muscle and liver, respectively in mice receiving X203 compared to those given IgG (**Fig. 3n**). As compared to 75 w/o mice, 100 w/o untreated and IgG-treated mice showed progressive telomere attrition and reduction in mtDNA copy number in liver, muscle and vWAT that was not seen in 100 w/o X203-treated mice (**Fig. 3o, p**).

### Specific beneficial effects of anti-IL11 in white adipose tissue

We performed bulk RNA-sequencing (RNA-seq) of vWAT, gastrocnemius, and liver isolated from IgG or anti-IL11-treated 100 w/o mice to further dissect molecular mechanisms. Across tissues, mice receiving anti-IL11 had the most significant gene set enrichment scores for hallmarks of metabolism, whereas markers of inflammation, EMT and cell cycle arrest were reduced (**Fig. 4a; Extended Data Table 5**).

**Figure 4.**
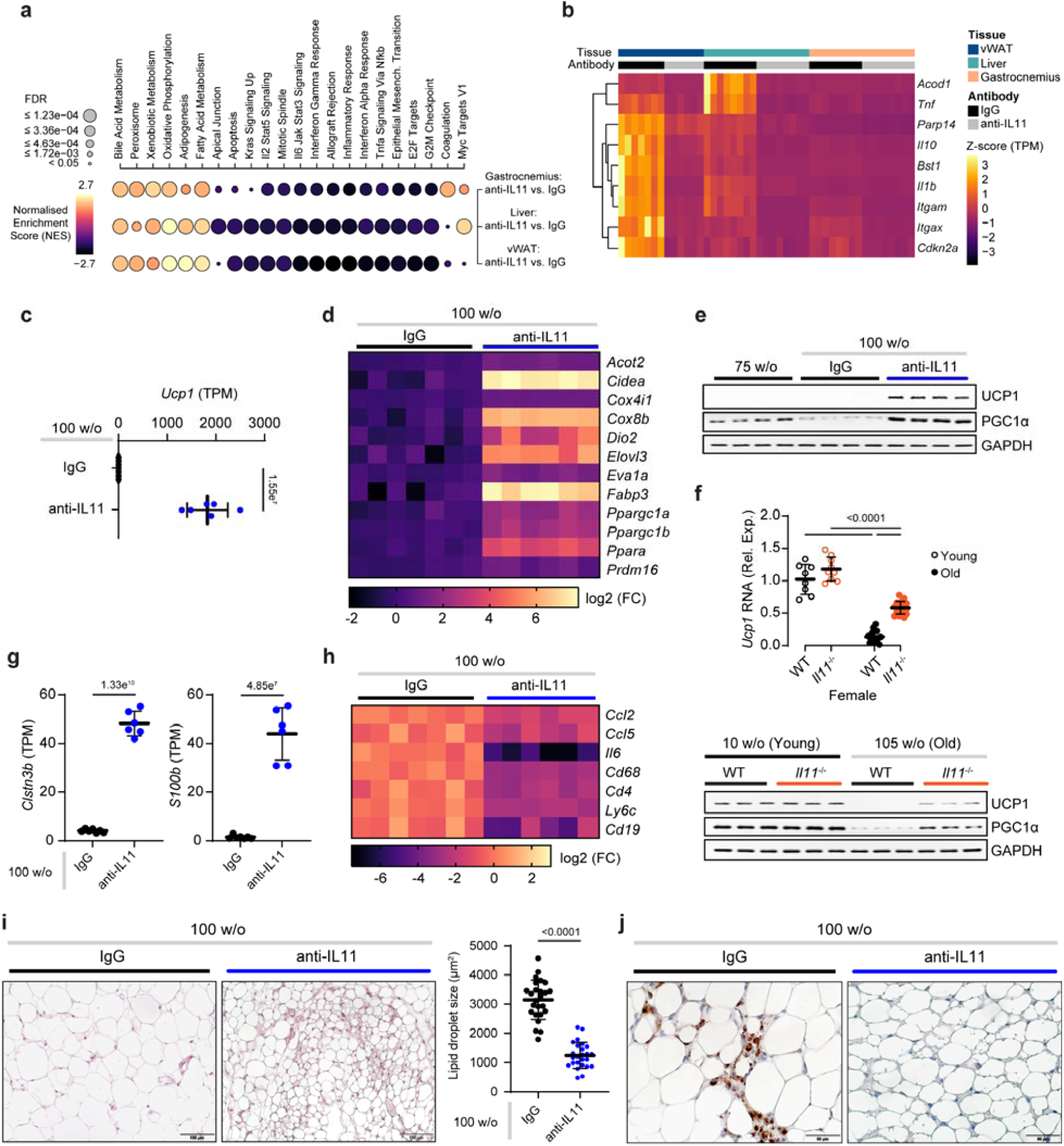
Anti-IL11 reduces inflammation and stimulates beiging in visceral white adipose tissue. **a-e, g-j** Data for X203 therapeutic experiments in old male mice as shown in Fig 3a. **a** Bubblemap showing results of hallmark gene set enrichment analysis for differentially expressed genes in the vWAT, liver, gastrocnemius of mice receiving anti-IL11 therapy as compared to IgG. Normalized enrichment scores (NES) are represented by colors, (black: negative NES, suggesting down-regulation of the gene set; yellow: positive NES, suggesting up-regulation). Dot size indicates significance (the larger the dot, the smaller the adjusted p-value). **b** Heatmap of row-wise scaled Transcripts per million (TPM) values of senescence genes in vWAT, liver, gastrocnemius, **c** abundance of *Ucp1* reads in vWAT (TPM), and **d** log2 FC heatmap of beijing genes in vWAT from IgG or anti-IL11 treated 100 w/o mice based on RNA-seq. **e** WB of Ucp1, Pgc1⍺ and GAPDH expression in vWAT (representative datasets from n=6/group). **f** Relative expression levels of *Ucp1* mRNA (young WT, n=8; young *Il11^-/-^*, n=9; old WT, n=16; old *Il11^-/-^*, n=18) as well as Ucp1 and Pgc⍺ protein rsin (representative datasets from n=6/group) in vWAT isolated from young and old female WT and *Il11^-/-^* mice. **g** Abundance of *Clstn3b* and *S100b* reads and **h** log2 FC heatmap of pro-inflammatory markers (from RNA-seq) in vWAT. **i** H&E-stained vWAT (scale bars, 100 µm) and quantification of lipid droplet size (mean of lipid droplet area), and **j** immunohistochemistry staining of CD68 in vWAT (scale bars, 50 µm). **a-d, f-h** Liver and gastrocnemius (n=8/group), vWAT IgG, n=7; vWAT anti-IL11, n=6. **c, f-g, i** Data are shown as meanLJ±LJSD. **c, g, i** Two-tailed Student’s t-test; **f** two-way ANOVA with Sidak’s correction.

The Tabula Muris Senis consortium identified a group of senescence genes (*Cdkn2a, Tnf, Il10, Il1b, Bst1, Irg1, Parp14, Itgax, Itgam*^28^) that are strongly and specifically upregulated in ageing that we used to study tissue-specific senescence effects. In vWAT, there was large upregulation of many genes in this gene signature, which were decreased by anti-IL11 (**Fig. 4b; Extended Data Fig. 7a**). The senescence gene signature was also present, albeit less so, in muscle and liver where X203 reduced its expression. As such, senescence and the effects of anti-IL11 on senescence markers were most apparent in vWAT.

### Anti-IL11 stimulates white fat beiging

More detailed study of the vWAT transcriptome in mice receiving X203 or IgG revealed that the gene most upregulated by anti-IL11 genome wide was *Ucp1* (**Fig. 4c; Extended Data Table 5**). *UCP1* is a canonical BAT gene and important for the development of thermogenic ‘beige/brite’ adipocytes in WAT deposits, which has large metabolic benefits^42, 43^. Of note, loss of thermogenic adipocytes in WAT is an ageing phenotype in mice and humans^8–10^.

On closer inspection, there was upregulation of a larger beiging program (*Acot2*, *Cidea*, *Cox4i1*, *Cox8b*, *Dio2*, *Elovl3*, *Eva1a*, *Fabp3*, *Ppargc1a*, *Ppargc1b*, *Ppara* and *Prdm16*) in vWAT of mice receiving anti-IL11 (**Fig. 4d**). Upregulation of UCP1 and PGC1α was validated at the protein level and further associated with *Il11* loss-of-function in old female *Il11^-/-^*mice (**Fig. 4e, f**). We also showed *Il11*-dependent, age-related suppression of *Ucp1* expression in vWAT of male and female mice deleted for *Il11ra1* (**Extended Data Fig. 7b**). PGC1α is a master regulator of mitochondrial biogenesis and we more specifically profiled expression of mitochondrial genes in vWAT (**Extended Data Fig. 7c**). Mitochondrial pathway analysis revealed significant increases in terms associated with enhanced mitochondrial biogenesis and function (**Extended Data Fig. 7d**). We did not extend molecular analyses to scWAT given the minimal amounts recovered from old *Il11^-/-^* mice and mice on anti-IL11 (**Fig. 2m**; **Extended Data Fig. 5l, 6g**).

In mice receiving X203, we also observed strong upregulation of *Clstn3b*, a newly identified mammal-specific product of the 3’ end of the *Clstn3* locus that promotes WAT triglyceride metabolism in partnership with *S100b*^44, 45^, which was also upregulated (**Fig. 4g; Extended Data Fig. 7e**). There was limited downregulation of *Ucp1* in BAT with age in WT mice and *Ucp1* was mildly elevated in BAT of *Il11^-/-^* mice but not in mice on anti-IL11 (**Extended Data Fig. 7f, g**).

These data likely explain why mice deleted for *Il11ra1* (**Fig. 1h**) or *Il11* (**Fig. 2f**) and mice administered anti-IL11 (**Fig. 3h**) have elevated body temperatures and are lean with favourable metabolic profiles even though they eat more than controls.

### Anti-IL11 reduces adipose inflammation and adipocyte size in old mice

The expression of pro-inflammatory genes (*Ccl2, Ccl5, Tnf*L*, Il6, Il1*β) in vWAT was higher in mice receiving IgG as compared to those on X203 (**Fig. 4b, h**), mirroring findings in livers of *Il11ra1^-/-^* mice and vWAT of *Il11^-/-^*mice (**Extended Data Fig. 2g, 4i**). We extended analyses to vWAT of young and old *Il11ra1^-/^*^-^ and WT mice of both sexes and confirmed age-dependent pro-inflammatory gene expression in WT mice, which was reduced in mice of *Il11ra1^-/^*^-^ genotype across sexes (**Extended Data Fig. 7h**).

Stromal inflammation is associated with immune cell infiltration and immune cell surface markers (*Cd68*, *Cd4*, *Ly6C*, and *Cd19*) were downregulated in the vWAT of mice receiving X203, as compared to IgG (**Fig. 4h**). Histology studies revealed that vWAT of X203-treated mice had an average 2.5-fold reduction in lipid droplet area, foci of beige adipocytes and fewer CD68^+ve^ macrophages, as compared to the IgG group (**Fig. 4i, j**).

### Engineering of an anti-IL11 antibody for clinical trials

Our data suggested IL11 inhibition using a therapeutic antibody approach as a potential strategy for extending human healthspan. Following on from our earlier finding that IL11 is a therapeutic target for pulmonary fibrosis^41^ there has been interest in IL11 inhibition to treat lung disease and clinical safety trials of intravenous anti-IL11RA (NCT05331300) or anti-IL11 (NCT05740475 and NCT05658107) have been initiated. However, long term neutralisation of IL11 in humans, as needed to extend healthspan, requires a highly efficacious molecule to facilitate subcutaneous drug administration and to limit dosing frequency. We thus engineered a neutralising IL11 antibody with these goals in mind.

High throughput screening of a panel of murine monoclonal antibodies for affinity and cross-species reactivity and IL11 neutralisation prioritised a lead clone VVB011, with a dissociation constant (K*_D_*) of 16 pM for human IL11 and strong binding in the mouse (60 pM) and cynomolgus monkey (94 pM) (**Fig. 5a**; **Extended Data Fig. 8a**). Robust neutralisation of species-matched IL11 signalling and bioactivity was observed across species, using an established MMP2 readout^46^ and immunoblotting (**Fig. 5b**; **Extended Data Fig. 8b**). The murine antibody had a half-life of >9 days (**Fig. 5c**), To optimise VVB011, we removed sequence liabilities and humanised the antibody that led to a further improvement in affinity into the single-digit picomolar range along with excellent neutralisation properties (**Fig. 5d, e**).

**Fig. 5.**
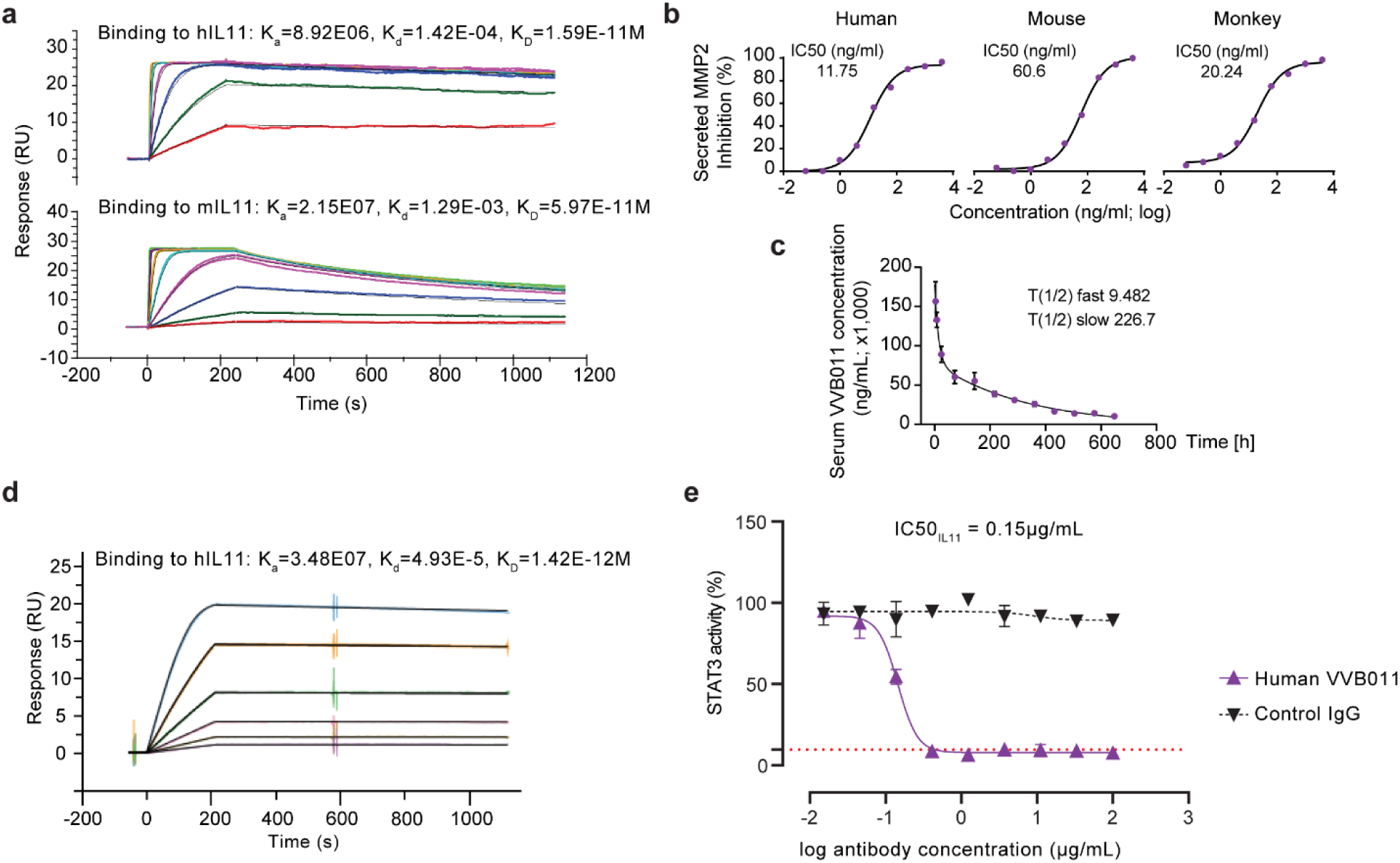
Development and engineering of a humanised anti-IL11 antibody (VVB011). **a** VVB011 interaction with human IL11 (top) or mouse IL11 (bottom) as determined by surface plasmon resonance (SPR). **b** Dose-response curve and half maximal inhibitory concentration value of VVB011 (61 pg/mL to 4 μg/mL) in inhibiting matrix metalloproteinase 2 (MMP2) secretion by hIL11-stimulated human cardiac fibroblasts, mIL11-stimulated mouse atrial fibroblasts, and cIL11-stimulated monkey dermal fibroblasts. **c** Blood pharmacokinetics of VVB011 in rodents (n=3). **d** Binding affinity and kinetics of humanised VVB011 to hIL11 by SPR. **e** STAT3-luciferase reporter activity of IL11-stimulated HEK293 in the presence of varying concentration of IgG control or humanised VVB011 clone (n=2). **c,e** Data are shown as meanLJ±LJSD.

## Discussion

Here we show that IL11 regulates the key ageing pathways to reduce healthspan. We propose that the many benefits seen with IL11 inhibition reflect its modulation across ageing pathways, as seen using polypharmacy in flies^7^. While IL11 is little studied and not previously thought important for healthspan, in retrospect, genomic data pointed to its involvement: SNPs at the *IL11* locus are associated with osteoarthritis^47^ and menopause^48^ and *Il11* is highly upregulated in idiopathic pulmonary fibrosis, common in the elderly and telomere disorders^49^. Recently, serum IL11 levels were shown to be elevated in the very old^50^.

The metabolic effects seen with inhibition of IL11 signalling phenocopy those of mice with WAT-specific deletion of Raptor that are lean with vWAT browning and we surmise that inhibition of IL11 prevents mTORC1 activation in old adipocytes/precursors, affording beiging^42, 43^. In a similar fashion, improved liver function likely reflects restored AMPK activity in hepatocytes^51^. Beiging of WAT has long been proposed as a therapeutic strategy to treat obesity and diabetes and anti-IL11 may represent a means of reactivating WAT beiging in the elderly.

Inhibition of IL11 had notable non-metabolic benefits, including reduced vestibular disturbance, vision loss, coat depigmentation and sarcopenia: suggestive of healthspan improvements at the level of the whole organism, across hallmarks and biomarkers of ageing. This begs the question as to whether inhibition of IL11 has beneficial effects in other aged tissues and/or on lifespan. In support of this notion, IL11 was newly suggested as a therapeutic target in osteoarthritis, a canonical disease of old age^47, 52^.

Inhibition of ERK or mTORC1 and activation of LKB1/AMPK by trametinib, rapamycin or metformin respectively increase lifespan in model organisms and are advocated by some for use in humans. However, these agents have on-target toxicities and can cause extended frailty, rather than improving healthspan^12, 13, 34^. We end by suggesting anti-IL11, which has a reassuring safety profile^41, 53^, as a new therapeutic approach for extending mammalian healthspan.

## Acknowledgements

This research is supported by the National Medical Research Council (NMRC)/STaR/0029/2017 (S.A.C.), NMRC Centre Grant to the NHCS (S.A.C.), MOHLJCIRG18novLJ0002 (S.A.C.), Tanoto Foundation (S.A.C.), Leducq Foundation (S.A.C.), NMRC/OFYIRG/0053/2017 (A.A.W.), NMRC MOH-OFIRG21nov-0006 (A.A.W.), Duke-NUS KBrFA/2022/0057 from Khoo Foundation (A.A.W.), and Goh Foundation (S.A.C. and A.A.W.). N.H is supported by Leducq Foundation 16CVD03, ERC advanced grant under the European Union Horizon 2020 Research and Innovation Program (AdG788970), and Deutsche Forschungsgemeinschaft (DFG-German Research Foundation) SFB 1470 HFpEF. W.W.L. is supported by A*STAR AME YIRG (A2084c0157). B.K.S. is supported by MOH-OFIRG19may-0002. CRUK (C15075/A28647) funds research in J.G’s laboratory. D.J.W., S.A.C., D.C. and J.G. are supported by the LMS institute of the MRC.

## Author contributions

A.A.W. and S.A.C. conceived, designed, funded, and provided supervision for the study. A.A.W., W.W.L., S.V., B.C., J.W.T.G, J.T., B.K.S., B.L.G., E.A., M.S., M.T. performed in vitro cell culture, in vivo studies, biochemistry, and molecular biology experiments. W.W.L., S.G.S., S.Y.L., and C.X. performed histology analysis. C.J.P. performed RNA-seq. R.L. and L.H performed phenomaster study. S.W. provided resources. A.A.W., W.W.L., S.C., C.J.P., R.L., B.K.S., D.J.W., N.H., S.S., L.H., J.G., D.C., and S.A.C. analysed the data. A.A.W., W.W.L., and S.A.C. prepared the manuscript with input from co-authors.

## Methods

### Antibodies

Commercial antibodies: Adiponectin (AdipoQ, 21613-1-AP, Proteintech), phospho-AMPK Thr172 (2535, CST), AMPK (5832, CST), CD31 (ab222783, clone EPR17260-263, abcam), CD68 (ab12512, abcam), Cyclin D1 (55506, clone E3P5S, CST), phospho-ERK1/2 Thr202/Tyr204 (4370, clone D13.14.4E, CST), ERK1/2 (4695, clone 137F5, CST), GAPDH (2118, clone 14C10, CST), FHL1 (10991-1-AP, Proteintech), GFP (ab290, abcam), phospho-LKB1 Ser428 (3482, clone C67A3, CST), LKB1 (3047S, clone D60C5, CST), phospho-mTOR Ser2448 (2971, CST), mTOR (2972,CST), p16 (human, ab108349, clone EPR1473, abcam), p16 (mouse, ab232402, clone EPR20418abcam), p21 (human, ab109520, clone EPR362 abcam), p21 (mouse, ab188224, clone EPR18021, abcam), phospho-p70S6K Thr389 (9234, clone 108D2, CST), p70S6K (2708, clone 49D7, CST), phospho-p90RSK Ser380 (11989, clone D3H11, CST), p90RSK (9355, clone 32D7, CST), PDGFR⍺ (AF1062, R&D systems), phospho-S6 ribosomal protein Ser235/236 (4858, CST), S6 ribosomal protein (2217, CST), PCNA (13110, clone D3H8P, CST), PGC1⍺ (ab191838, abcam) SLC10A1 (MBS177905, MyBioSource), SM22⍺ (ab14106, abcam), phospho-STAT3 Tyr705 (4113, clone M9C6, CST), STAT3 (4904, clone 79D7, CST), UCP1 (72298, clone E9Z2V, CST), anti-rabbit HRP (7074, CST), anti-mouse HRP (7076, CST), anti-rabbit Alexa Fluor 488 (ab150077, abcam), anti-goat Alexa Fluor 488 (ab150129, abcam), anti-rabbit Alexa Fluor 555 (ab150074, abcam). All commercially available antibodies have been validated by their manufacturer as indicated in their respective datasheet and/or website.

Custom-made antibodies: IgG (clone 11E10), anti-IL11 (clone X203 for both WB and neutralising studies), humanised anti-IL11 (clone VVB011 for neutralising study), anti-IL11RA (clone X209 for neutralising study) were manufactured by Genovac. IgG (11E10) suitability as a control antibody was validated previously^22^. X203 was validated for neutralisation of human and mouse IL11^22, 46^, and for WB^23, 46^. X209 was validated previously for neutralisation of human and mouse IL11RA^46^ and for WB^46^. VVB011 was validated for binding to and neutralising human, mouse, and cynomolgus IL11 (in this manuscript).

### Recombinant proteins

Recombinant cynomolgus IL 11 (cIL11, 90925-CNCE, Sino Biological), human IL11 (hIL11, Z03108, Genscript), mouse IL11 (mIL11, Z03052, Genscript).

### Chemicals

Bovine serum albumin (BSA, A7906, Sigma), 16% Formaldehyde (w/v), methanol-free (28908, Thermo Fisher Scientific), DAPI (D1306, Thermo Fisher Scientific), DMSO (D2650, Sigma), Rapamycin, (9904, CST), Triton X-100 (T8787, Sigma), Tween-20 (170-6531, Bio-Rad), U0126 (9903, CST),

### Ethics statements

All experimental protocols involving human subjects (commercial primary human cell lines) were performed in accordance with the *ICH Guidelines for Good Clinical Practice*. All participants provided written informed consent and ethical approvals have been obtained by the relevant parties as written in the datasheets provided by either ScienCell and ATCC from which primary human cardiac fibroblasts, primary human hepatocytes, and A549 were commercially sourced.

Animal studies were carried out in compliance with the recommendations in the Guidelines on the Care and Use of Animals for Scientific Purposes of the National Advisory Committee for Laboratory Animal Research (NACLAR). All experimental procedures were approved (SHS/2019/1481 and SHS/2019/1483) and conducted in accordance with the SingHealth Institutional Animal Care and Use Committee (IACUC).

### Cell culture

Cells were grown and maintained at 37 °C and 5% CO2.The growth medium was renewed every 2–3 days and cells were passaged at 80% confluence, using standard trypsinization techniques. All experiments were carried out at P3, unless otherwise specified. Cells were serum-starved overnight in basal media prior to stimulation with different treatment conditions (in the absence or presence of antibodies or inhibitors) and durations, as outlined in the main text or figure legends.

1. Primary human cardiac fibroblasts. Primary human cardiac fibroblasts (HCFs, 52-year-old male, 6330, ScienCell) were grown and maintained in complete fibroblasts medium-2 (2331, ScienCell) supplemented with 5% foetal bovine serum (FBS, 0500, ScienCell), 1% fibroblasts growth supplement-2 (FGS-2, 2382, ScienCell) and 1% penicillin-streptomycin (P/S, 0513, ScienCell). For replicative senescence study, primary HCFs were serially passaged (from passage 4 (P4) to passage 14 (P14)) in the absence or presence of a neutralising IL11RA antibody (X209) or an IgG isotype control (11E10).
2. Primary human hepatocytes. Primary human hepatocytes were isolated from a 22-week-old foetus (5200, ScienCell). Following recovery from the initial thaw cycle, hepatocytes were seeded at a density 4X10^5^ cells/well of a collagen-coated 6-well plate and maintained in hepatocyte medium (5201, ScienCell) which contains 2% FBS and 1% P/S. Hepatocytes were then used directly for downstream experiments within 48 hours of seeding.
3. A549. Human lung epithelial carcinoma cells (A549 (58-year-old male), ATCC) were grown and maintained in DMEM (11995065, Gibco) complete medium which contains 10% FBS (10500064, Gibco) and 1% P/S (15140122, Gibco).
4. Primary monkey dermal fibroblasts. Primary monkey dermal fibroblasts (P10843, Innoprot) were maintained in fibroblast complete medium (P60108, Innoprot).
5. Primary mouse atrial fibroblasts Primary mouse atrial fibroblasts (MAFs) were isolated from wild-type of B6.129S1-*Il11ra^tm1Wehi^*/J mouse strain (The Jackson Laboratory). Mouse atria were minced and digested with mild agitation for 30 minutes at 37°C in DMEM containing 1% P/S and 0.14 Wunsch U ml^-1^ Liberase (5401119001, Roche). MAFs were enriched via negative selection with magnetic beads against mouse CD45 (leukocytes), CD31 (endothelial) and CD326 (epithelial) using a QuadroMACS separator (Miltenyi Biotec) according to the manufacturer’s protocol. MAFs were then maintained in complete DMEM supplemented with 10% FBS and 1% P/S.

### Olink proximity extension assay

Human hepatocytes were seeded at a density of 2.5X10^5^ cells/well into 6-well plates. The culture supernatants were collected following stimulation with IL11 (0, 6, and 24 hours) and were sent to the Olink Proteomics, Uppsala, Sweden for proximity extension assays using the 92-protein inflammation panel. The protein concentrations were expressed as normalised protein expression (NPX; log2 scale) and those proteins with concentrations below the limit of detection were excluded from analysis.

### Operetta high throughput phenotyping assay

HCFs (P4, P7, P10, and P14) were seeded in 96-well black CellCarrier plates (PerkinElmer) at a density of 6x10^3^ cells/well either untreated or in the presence of IgG or X209. After reaching ∼80% confluence, cells were fixed in 4% formaldehyde, and permeabilized with 0.1% Triton X-100. Non-specific sites were blocked with blocking solutions (0.5% BSA and 0.1% Tween-20 in PBS). Cells were incubated overnight (4°C) with primary antibodies (p16 and p21) at a dilution of 1:500, followed by incubation with the appropriate Alexa Fluor 488 secondary antibodies (1:1000, 1 hour, RT). Cells were then counterstained with 1 µg/ml DAPI in blocking solution. Antibodies and DAPI were diluted in blocking solutions. Each condition was imaged from duplicated wells and a minimum of 7 fields/well using Operetta high-content imaging system 1483 (PerkinElmer). The measurement of p16 and p21 fluorescence intensity per area (normalised to the number of cells) was performed with Columbus 2.9 (PerkinElmer).

### Seahorse assay

Primary HCFs were seeded into the Seahorse XF 96-well Cell Culture Microplate (40x10^3^ cells/well) and serum-starved overnight prior to stimulations. Seahorse measurements were performed on Seahorse XFe96 Extracellular Flux analyzer (Agilent). XF Cell Mito Stress Test kit (103015-100, Agilent) and Seahorse XF Mito Fuel Flex Test kit (103260-100, Agilent) were used according to the manufacturer’s protocol to measure the mitochondrial oxygen consumption rate and the percentage of fatty acid oxidation, respectively as described previously^24^. Seahorse Wave Desktop software (Ver 2.6.3) was used for report generation and data analysis.

### Surface plasmon resonance assay

Binding kinetics of VVB011 (ligand) to analytes i.e. human, mouse, and cynomolgus IL11 were measured on Biacore T200 (GE Healthcare). VVB011 was immobilised on an anti-mouse capture chips and a concentration range of each of the analytes (hIL11: 0.37 nM to 90 nM; mIL11: 0.04nM to 90nM; cIL11: 0.23nM to 30nM) was injected at a flow rate of 30 μl/min (hIL11), 40 μl/min (mIL11) or 50 μl/min (cIL11). Interaction assays were performed with HBS-P+ and 1 mg/ml BSA. Affinity and kinetic constants were determined by fitting the corrected sensograms assuming 1:1 binding with double reference subtraction. The equilibrium binding constant *K*_D_ was determined by the ratio of *k*d/*k*a.

### Half Maximal Inhibitory Concentration Measurement

HCFs, MAFs, and monkey dermal fibroblasts were stimulated with IL11 (24 hours) in the presence of IgG (11E10, 2 µg/ml) or VVB011 (61 pg/ml to 4 µg/ml; 4-fold dilutions).

Supernatants were collected and assayed for the amount of secreted MMP2 (MMP200, R&D Systems). Dose-response curves were generated by plotting the logarithm of VVB011 concentrations versus the corresponding inhibition values (%), using the least squares ordinary fit. We considered MMP2 secretion by unstimulated cells as maximal inhibition (100%), while MMP2 secretion after stimulation with IL11 in the presence of 11E10 constituted 0% inhibition.

### Animal models

All mice were housed at 21-24L with 40-70% humidity on a 12-hour light/dark cycle and provided food and water *ad libitum*.

1. *Il11ra1*-deleted mice (*Il11ra1*^−/−^ or *Il11ra1* KO) Male and female *Il11ra1^+/+^* (wild-type) and *Il11ra1^-/-^* mice^21^ (B6.129S1-Il11ra^tm1Wehi^/J, The Jackson Laboratory) were sacrificed at 110 weeks of age for blood and tissue collection; 10-12 weeks old male and female mice of the respective genotypes were used as controls.
2. *Il11*-deleted mice (*Il11*^−/−^) Male and female mice lacking functional alleles for *Il11* (*Il11*^−/−^), which were generated and characterised previously^23, 32^, and their wild-type counterparts were sacrificed at 10-12 weeks of age (young controls) and 104-108 weeks of age (old group).
3. *Il11-EGFP* reporter mice Young (10-week-old) and old (100-week-old) transgenic mice (C57BL/6J background) with *EGFP* constitutively knocked-into the *Il11* gene (*Il11-EGFP* mice, Cyagen Biosciences Inc)^22^ were sacrificed for immunofluorescence staining studies on liver, gastrocnemius, and visceral gonadal white adipose tissue (referred in the text as vWAT). Old wild-type littermates were used as negative controls.
4. In vivo administration of anti-IL11 Male C57Bl/6J mice (Jackson Laboratory) were randomised prior to receiving either no treatment, or anti-IL11 (X203 or VVB011) or IgG (11E10) treatment. X203 or 11E10 (40 mg/kg, every 3 weeks) were administered by intraperitoneal (IP) injection, starting from 75 weeks of age for a duration of 25 weeks; mice were then sacrificed at 100 weeks of age. For VVB011 *in vivo* study, 85-week-old mice were administered either VVB011 or 11E10 (40 mg/kg, every 2 weeks) for 10 weeks.

### Glucose tolerance test (GTT) and insulin tolerance test (ITT)

Mice were fasted for 6 hours prior to baseline blood glucose measurement. For GTT, mice were injected intraperitoneally with 20% glucose at 2 mg/g lean mass. For ITT, mice were injected intraperitoneally with recombinant human insulin at 1.2 mU/g body weight. Both glucose and insulin were diluted in sterile DPBS. Blood glucose concentrations were then measured at 15, 30, 60, 75, and 90 minutes after glucose or insulin administration, for GTT or ITT, respectively. Blood was collected via tail snip and Accu-Chek blood glucometer was used for blood glucose measurements.

### Echo MRI

Mouse body composition (total body fat and lean mass measurements) was performed 1 day prior to GTT/ITT or sacrifice by EchoMRI analysis using 4in1 Composition Analyzer for live small animals (Echo Medical Systems).

### Frailty scoring

End-point frailty scoring was performed, with observers blinded to treatment, 1-2 days prior to sacrifice using a 30-point frailty scoring system^33^. Body temperatures were recorded by means of rectal thermometry using Kimo Thermocouple Thermometer (TK110, Kimo).

### Grip strength assessment

A digital grip strength meter (BIO-GS3, BIOSEB) was used to measure full body (4 limbs) and forelimb (both forepaws) grip strengths, as per the manufacturer’s instruction. Mice were allowed to rest for at least 1 hour between the two tests. The average of 3 readings of maximal average force exerted by each mouse on the grip strength meter was used for analysis.

### Measurement of whole-body metabolic parameters (PhenoMaster)

Whole-body metabolic parameters for IgG/X203-treated (Ab cohort) and wild-type/IL11 KO (KO cohort) mice were assessed by open-circuit indirect calorimetry. Animals were single-housed in the PhenoMaster automated home-cage system (TSE Systems) in a temperature (22L) and humidity-controlled environment with a 12-hour light/dark cycle. Parameters including oxygen consumption (VO2), carbon dioxide production (VCO2), food intake, and locomotor activity were measured simultaneously at 1 minute time intervals. Respiratory exchange ratio (RER) was calculated using the VCO2/VO2 ratio. Locomotor activity was divided into horizontal plane locomotor activities, defined as the total number of infrared beam breaks in the X- and Y-axis (counts). Mice were monitored for 5 consecutive overnight periods including an acclimatisation period during the first light/dark cycle (day 0-1) which was not used for analysis. For both Ab and KO cohorts, the control (IgG/WT) group (n=10) and intervention (X203/IL11 KO) group (n=10) were divided equally into two consecutive monitoring sessions where animals from each group (n=5) were allocated alternating PhenoMaster cage locations that were swapped in the second session. Young (14-week-old) wild-type animals (n=10) used for baseline RER comparison in the Ab cohort were monitored separately in two sessions (n=5 per session). Baseline RER comparison was made using measurements from the second light/dark cycle (day 1-2). Animals were given *ad libitum* access to food and water except during test phases introduced after day 2 where food access was restricted to assess the resting metabolic rate (measured at thermoneutrality (28L)) and adaptation to fasting (12 hours).

### Epigenetic ageing clock measurement (DNAge®)

The measurements of the epigenetic ageing clock were performed using the DNAge® Service (Zymo Research). Briefly, DNA extraction and clean-up were performed followed by the bisulfite conversion process. Samples were enriched specifically for the sequencing of >1000 age-associated gene loci using Simplified Whole-panel Amplification Reaction Method (SWARM®), where specific CpGs are sequenced at minimum 1000X coverage. Sequencing was run on an Illumina NovaSeq instrument. Sequences were identified by Illumina base calling software then aligned to the reference genome using Bismark. Methylation levels for each cytosine were calculated by dividing the number of reads reporting a “c” by the number of reads reporting a “C” or “T”. The percentage of methylation for these specific sequences were used to assess DNA age according to Zymo Research’s proprietary DNAge® predictor which had been established using elastic net regression to determine the DNAge®.

### Pharmacokinetic study

The pharmacokinetic study of VVB011 was outsourced to WuXi Apptec. Briefly, VVB011 was injected into rodents (n=3) with a dose of 5 mg/kg by intravenous bolus. At timepoints following injection, the concentration of VVB011 in the serum was determined using a human IL11 ELISA (assay range was 1000-16000 ng/ml). The signal was measured using a SpectraMax M5e/M5/Plus 384 plate reader (Molecular Devices) and the curve was fitted using a 4-Parameter Logistic and weighting factor 1/Y^2^. To estimate the half-life of both the initial distribution (T1/2 fast) and the elimination phase (T1/2 slow), data were fit using nonlinear regression - two phase exponential decay.

### Colorimetric and enzyme-linked immunosorbent assays (ELISA)

The levels of alanine transaminase (ALT), aspartate aminotransferase (AST), cholesterol, and β-hydroxybutyrate (Ketone body), and IL6 in mouse serum were measured using Alanine Transaminase Activity Assay Kit (ab105134, abcam), Aspartate Aminotransferase Activity Assay Kit (ab105135, abcam), Cholesterol Assay Kit (ab65390, Abcam), beta-hydroxybutyrate Colorimetric Assay Kit (700190; Cayman chemicals), and Mouse IL-6 Quantikine ELISA Kit (M6000B, R&D Systems), respectively. The levels of triglyceride in mouse livers and serum were measured using Triglyceride Assay Kit (ab65336, Abcam). Total collagen content in mouse livers, gastrocnemius, and vWAT were measured using Quickzyme Total Collagen assay kit (QZBtotco15, Quickzyme Biosciences). The levels of IL6, IL8, and IL11 in equal volumes of cell culture media collected from experiments with primary human cells were quantified using Human IL-8/CXCL8 Quantikine ELISA Kit (D8000C, R&D Systems), Human IL-6 Quantikine ELISA Kit (D6050, R&D Systems), Human IL11 Quantikine ELISA kit (D1100, R&D Systems). All ELISA and colorimetric assays were performed according to the manufacturer’s protocol.

### Triglyceride Assay Kit (ab65336, Abcam), Immunoblotting

Western blots were carried out on total protein extracts from liver, gastrocnemius, and vWAT tissues, which were homogenised in RIPA Lysis and Extraction Buffer (89901, Thermo Fisher Scientific) containing protease and phosphatase inhibitors (A32965 and A32957, Thermo Fisher Scientific). Protein lysates were separated by SDS-PAGE, transferred to PVDF membranes, blocked for 1 hour with 3% BSA, and incubated overnight with primary antibodies (1:1000). Protein bands were visualised using the SuperSignal^TM^ West Femto Maximum Sensitivity Substrate detection system (34096, Thermo Fisher Scientific) with the appropriate HRP secondary antibodies (1:5000).

### RT-qPCR

Total RNA was extracted from cells or snap-frozen tissues using TRIzol™ Reagent (15596026, Thermo Fisher Scientific) and RNeasy Mini Kit (74104, Qiagen). PCR amplifications were performed using iScript cDNA Synthesis Kit (1708891, Bio Rad). Gene expression analysis was performed with QuantiNova SYBR Green PCR Kit (208056, Qiagen) technology using StepOnePlus^TM^ (Applied Biosystem). Expression data were normalised to *GAPDH* mRNA expression and fold change was calculated using 2^-ΔΔCt^ method. The primer sequences are provided in Supplementary Table 1.

### Telomere length and mitochondrial copy number quantification

DNA from HCFs (P4 and P14) and snap-frozen liver, gastrocnemius, and vWAT was extracted with the E.Z.N.A. Ⓡ Tissue DNA Kit (D3396-02, Omega Bio-tek) according to the manufacturer’s protocol. Telomere length and mitochondrial copy number for HCFs were evaluated by RT-qPCR with the Relative Human Telomere Length Quantification qPCR Assay Kit (8908, ScienCell) and Relative Human Mitochondrial DNA copy number Length Quantification qPCR Assay Kit (8938, ScienCell), respectively. Similarly, the telomere length and mitochondrial copy number for mouse tissues were evaluated by RT-qPCR with the Relative Mouse Telomere Length Quantification qPCR Assay Kit (M8908, ScienCell) and Relative Human Mitochondrial DNA copy number Length Quantification qPCR Assay Kit (M8938, ScienCell), respectively.

### Histology

Investigators performing histology and analysis were blinded to the genotype/treatment group.

1. Hematoxylin and eosin (H&E) staining Mouse vWAT were fixed in 10% neutral-buffered formalin (NBF) for 48 hours, embedded in paraffin, cut into 4 μm sections followed by H&E staining according to the standard protocol. Lipid droplet areas were quantified by ImageJ (version 1.53t, NIH) with the adipocytes tools plugin (https://github.com/MontpellierRessourcesImagerie/imagej_macros_and_scripts/wiki/Adipocytes-Tools) from 5 randomly selected fields at 200X magnification in vWAT images per mouse, and 5 mice/groups were assessed. The mean value of lipid droplet areas per field was plotted for the final data presentation.
2. Immunohistochemistry (IHC) 4 μm mouse vWAT sections were dewaxed with histoclear and a gradient ethanol wash, followed by permeabilisation using 1% Triton-X 100 for 10 mins and antigen retrieval process with Reveal Decloaker (RV1000M, Biocare Medical) using a double boiler method at 110°C for 20 mins. Slides were allowed to cool in the container together with the Reveal Decloaker solution for 10 mins under running water. Double blocking was achieved with (1) H_2_O_2_ for 10 mins and (2) 2.5% normal horse serum for 1 hour (S-2012, Vector Labs). vWAT sections were incubated overnight at 4°C with primary antibody (CD68, 1:100 in PBST) and visualised by probing with Horse Anti-Rabbit IgG Polymer Kit (MP-7401, Vector Labs) for 1 hour at 37°C and ImmPACT DAB Peroxidase Substrate Kit (SK-4105, Vector Labs). Hematoxylin (H-3401, Vector Labs) was used to counterstain the nuclei prior to imaging by light microscopy (Olympus IX73).
3. Immunofluorescence Young (10-week) and aged (100-week) *Il11*^EGFP/+^ and aged wild-type (WT) *Il11*^+/+^ mice underwent perfusion-fixation with PBS and 4% PFA for multi-organ harvest at terminal sacrifice. Mouse liver, vWAT and gastrocnemius were further fixed in 4% PFA at 4°C and serial 15-30% sucrose dehydration over 48 hours before they were cryo-embedded in OCT medium. 5 µm sections were heat antigen retrieved using Reveal Decloaker (RV1000M, Biocare), permeabilized with 0.5% Triton X-100, and blocked with 5% normal horse serum before probing with primary antibodies at 4°C overnight. Alexa fluor conjugated secondary antibodies were incubated for 2 hours at room temperature for visualisation. Autofluorescence was quenched with 0.1% Sudan Black B for 20 minutes. DAPI was included for nuclear staining before mounting and sealed. Photomicrographs were randomly captured by researchers blinded to the strain and age groups.

### RNA sequencing (RNAseq) libraries

Total RNA was isolated from liver, fat and skeletal muscle of mice receiving either IgG or X203 using RNeasy Mini Kit (74104, Qiagen) and quantified using Qubit™ RNA Broad Range Assay Kit (Q10210, Thermo Fisher Scientific). RNA Quality Scores (RQS) were assessed using the RNA Assay (CLS960010, PerkinElmer) and DNA 5K/RNA/CZE HT Chip (760435, PerkinElmer) on a LabChip GX Touch HT Nucleic Acid Analyzer (CLS137031, PerkinElmer). TruSeq Stranded mRNA Library Prep kit (20020594, Illumina) was used to assess transcript abundance following the manufacturer’s instructions. Briefly, poly(A)+ RNA was purified from 1 µg of total RNA with RQS > 6, fragmented, and used for cDNA conversion, followed by 3’ adenylation, adaptor ligation, and PCR amplification. The final libraries were quantified using Qubit™ DNA Broad Range Assay Kit (Q32853, Thermo Fisher Scientific) according to the manufacturer’s guide. The average fragment size of the final libraries was determined using DNA 1K/12K/Hi Sensitivity Assay LabChip (760517, PerkinElmer)) and DNA High Sensitivity Reagent Kit (CLS760672, PerkinElmer). Libraries with unique dual indexes were pooled and sequenced on partial lanes targeting ∼50M reads per sample on a HiSeq or a NovaSeq 6000 sequencer (Illumina) using 150-bp paired-end sequencing chemistry.

### Data processing and analysis for RNAseq

Fastq files were generated by demultiplexing raw sequencing files (.bcl) with Illumina’s bcl2fastq v2.20.0.422 with the --no-lane-splitting option. Low quality read removal and adaptor trimming was carried out using Trimmomatic V0.36 with the options ILLUMINACLIP: <KEEPBOTHREADS>=TRUE MAXINFO:35:0.5 MINLEN:35. Reads were mapped to theMus musculus GRCm39 using STAR v.2.7.9a with the options --outFilterType BySJout -- outFilterMultimapNmax 20 --alignSJoverhangMin 8 --alignSJDBoverhangMin 1 -- outFilterMismatchNmax 999 --alignIntronMin 20 --alignIntronMax 1000000 --alignMatesGapMax 1000000 in paired end, single pass mode. Read counting at the gene-level was carried out using subread v.2.0.3: -t exon -g gene_id -O -s 2 -J -p -R -G. The Ensembl release 104 Mus musculus GRCm39 GTF was used as annotation to prepare STAR indexes and for FeatureCounts. Principal component analysis clustered samples into tissue-types and conditions. Outlier samples that did not cluster with the expected group were removed.

Differential expression (DE) genes were identified using R v4.2.0 using the Bioconductor package DESeq2 v1.36.0 using the Wald test for comparisons. IgG samples were used as the reference level for comparison with anti-IL11 (X203) samples for vWAT, liver, and gastrocnemius. Mitocarta v3.0 gene list was downloaded and TPM values in Fat IgG and anti- IL11 samples were plotted using pheatmap R package for genes which had TPM >=5 in at least one condition. Gene set enrichment analysis was carried out using the fgsea v.1.22.0 R package for MSigDB Hallmark (msigdbr v.7.5.1) and MitoCarta v3.0 gene sets with 100,000 iterations. The “stat” value quantified by DESeq2 was used to rank the genes, as an input for the enrichment analysis.

### Statistical analysis

Statistical analyses were performed using GraphPad Prism software (version 9.4.1). Datasets were tested for normality with Shapiro-Wilk tests. For normally distributed data, two-tailed Student’s *t*-tests or one-way ANOVA were used for analysing experimental setups requiring testing of two conditions or more than 2 conditions, respectively. P values were corrected for multiple testing according to Dunnett’s (when several experimental groups were compared to a single control group) or Tukey (when several conditions were compared to each other within one experiment). Comparison analysis for two parameters from two different groups were performed by two-way ANOVA and corrected with Sidak’s multiple comparisons. The Z-test is used when comparing the difference in the proportion of enlarged seminal vesicles between 2 groups.

The criterion for statistical significance was set at *P*□<□0.05.

### Data availability

All data are available within the Article or Supplementary Information. The RNAseq data reported in this paper are available on the Short Read Archive with Bioproject ID: PRJNA939262 (https://dataview.ncbi.nlm.nih.gov/object/PRJNA939262?reviewer=uv83svp0v6sfud6a0gboeuu7ub).

---

**Supplementary Fig 1.**
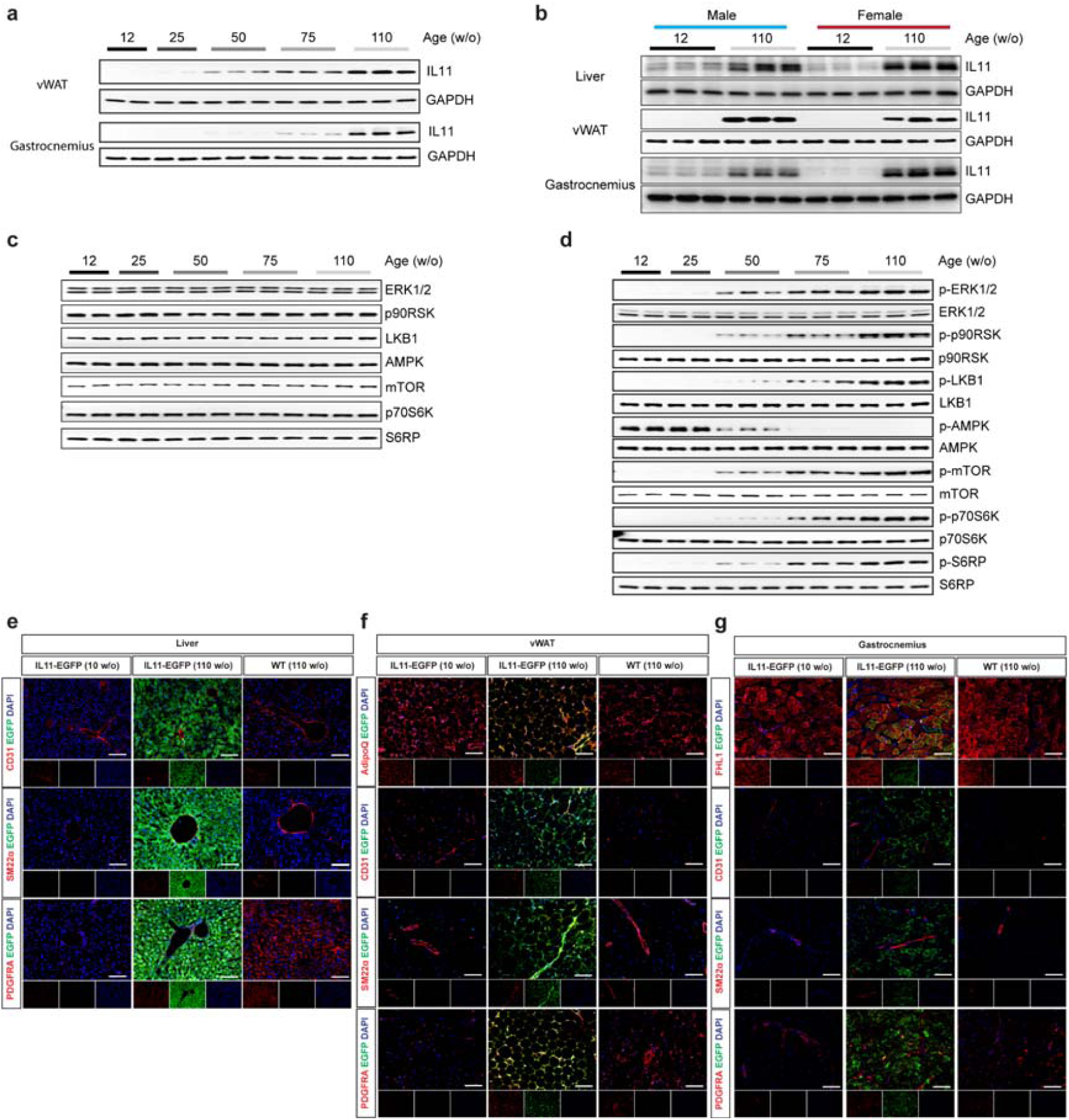
Age-dependent upregulation of IL11 in varied cell types across metabolic tissues. **a** Western blots (WB) of IL11 and GAPDH in visceral gonadal white adipose tissue (vWAT) and gastrocnemius from 12, 25, 50, 75, and 110-week-old (w/o) male mice (n=5/group). **b** WB of IL11 and GAPDH in the liver, vWAT and gastrocnemius from 12, and 110-week-old (w/o) male and female mice (n=3/group). **c** WB of ERK1/2, p90RSK, LKB1, AMPK, mTOR, p70S6K, and S6RP in livers from 12, 25, 50, 75, and 110-week-old (w/o) male mice (n=5/group) for the respective phopsho proteins shown in **Fig. 1b**. **d** WB of p-ERK1/2, p-p90RSK, p-LKB1, p-AMPK, p-mTOR, p-p70S6K, p-S6RP, and their respective total protein in gastrocnemius from 12, 25, 50, 75, and 110 w/o male mice (n=5/group). **e-g** Representative immunofluorescence images (scale bars, 100µm) of EGFP expression in the livers, vWAT, and gastrocnemius, colocalized with parenchymal cell markers Adiponectin (AdipoQ) in vWAT and Four and a half LIM domains (FHL1) in gastrocnemius, endothelial cells (CD31), smooth muscle transgelin (SM22lJ), and pan-fibroblast marker (PDGFRA) of 10 and 110-week old *Il11*-*EGFP* mice (representative dataset from n=3/group).

**Supplementary Fig 2.**
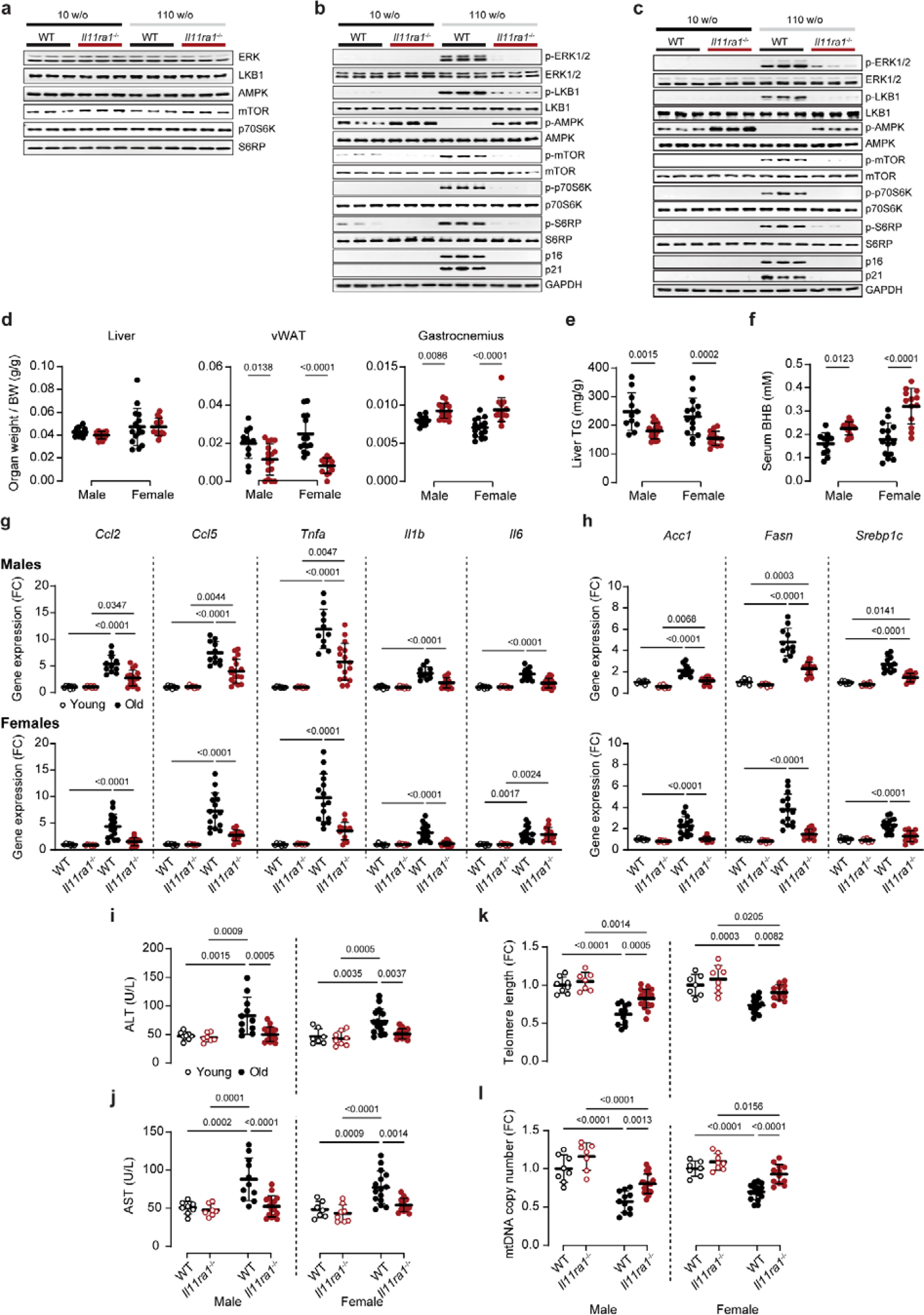
Beneficial signalling, metabolic, inflammation and ageing biomarker effects of *Il11ra1* deletion. **a** WB of total proteins for **Fig. 1e** (n=3/group). WB showing the activation status of ERK1/2, p90RSK, LKB1, AMPK, mTOR, p70S6K, S6RP, and protein expression levels of p16, p21 and GAPDH in **b** vWAT and **c** gastrocnemius from 10 and 110 w/o male WT and *Il11ra1^-/-^* mice (n=3/group). **d** Indexed weights of liver, vWAT and gastrocnemius, **e** liver triglycerides (TG) levels, **f** serum β-hydroxybutyrate (BHB) in 110 w/o male and female WT and *Il11ra1^-/-^* mice. Relative gene expression levels of **g** *Ccl2*, *Ccl5*, *Tnf*α, Il1flJ, *Il6*, **h** *Acc*, *Fasn* and *Srebp1c,* and serum levels of **i** AST and **j** ALT in young and old male and female WT and *Il11ra1^-/-^* mice. Gastrocnemius **k** telomere length and **l** mitochondria DNA (mtDNA) copy number from young and old male and female WT and *Il11ra1^-/-^* mice. **d-l** Data are shown as meanL±LSD (young male WT, n=8; young male *Il11ra1^-/-^*, n=7; old male WT, n=11-12; old male *Il11ra1^-/-^*, n=15-17; young female WT, n=7; young female *Il11ra1^-/-^*, n=8; old female WT, n=14-15; old female *Il11ra1^-/-^*, n=13); two-way ANOVA with Sidak’s correction.

**Supplementary Fig 3.**
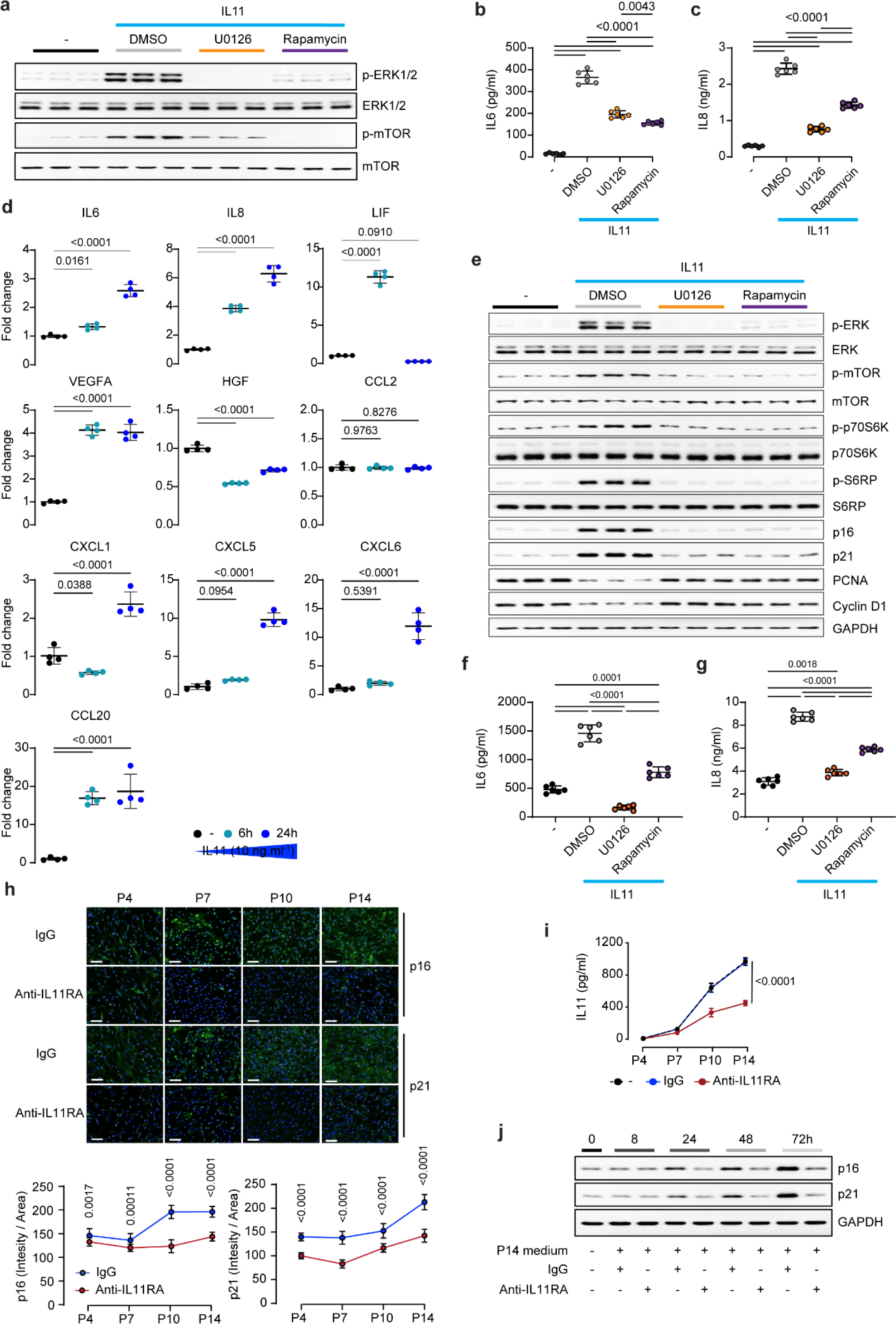
IL11 drives ERK/mTORC1-dependent senescence and senescence-associated secretory phenotypes. Effects of U0126 and rapamycin on **a** the activation status of ERK1/2 and mTOR by WB, and on the levels of secreted **b** IL6 and **c** IL8 by ELISA (n=6/group) from IL11 (5 ng/ml)-stimulated primary human cardiac fibroblasts. **d** Relative levels of IL6, IL8, LIF, VEGFA, HGF, CCL2, CXCL1, CXCL5, CXCL6, and CCL20 in the supernatant of IL11-stimulated primary human hepatocytes (6 and 24 hours) as measured by Olink proximity extension assay (n=4/group). **e-g** Data for IL11 (10 ng/ml)-stimulated primary human hepatocytes in the presence of either DMSO, U0126, or rapamycin (n=6/group). **e** WB showing the activation status of ERK1/2, mTOR, p70S6K, S6RP, and the protein expression levels p16, p21, PCNA, Cyclin D, and GAPDH. Concentrations of **f** IL6 and **g** IL8 in the supernatant (as measured by ELISA). **a-c, e-g** U0126 (10 µM), rapamycin (10 nM). **h** Immunofluorescence images (scale bars, 100 μm; representative datasets from n=7/group) and quantification of intensity/area (n=14/group) for p16 and p21 staining, and **i** concentrations of IL11 in the supernatant (by ELISA) of HCF passage 4 (P4), 7, 10, and 14 that had been passaged in the presence of either IgG or anti-IL11RA (X209; 2µg/ml) from P2. **j** WB showing the expression levels of p16, p21, and GAPDH from HCFs P4 that were stimulated for 8, 24, 48, and 72 hours with media collected from HCFs P14 that had been grown and passaged in the presence of either IgG or anti-IL11RA (X209; 2µg/ml) from P2 (representative datasets from n=4/group). **b-d, f-i** Data are shown as meanL±LSD. **b, c, f, g** one-way ANOVA with Tukey’s correction; **d** one-way ANOVA with Dunnett’s correction, **h** two-way ANOVA with Sidak’s correction, **i** two-way ANOVA.

**Supplementary Fig 4.**
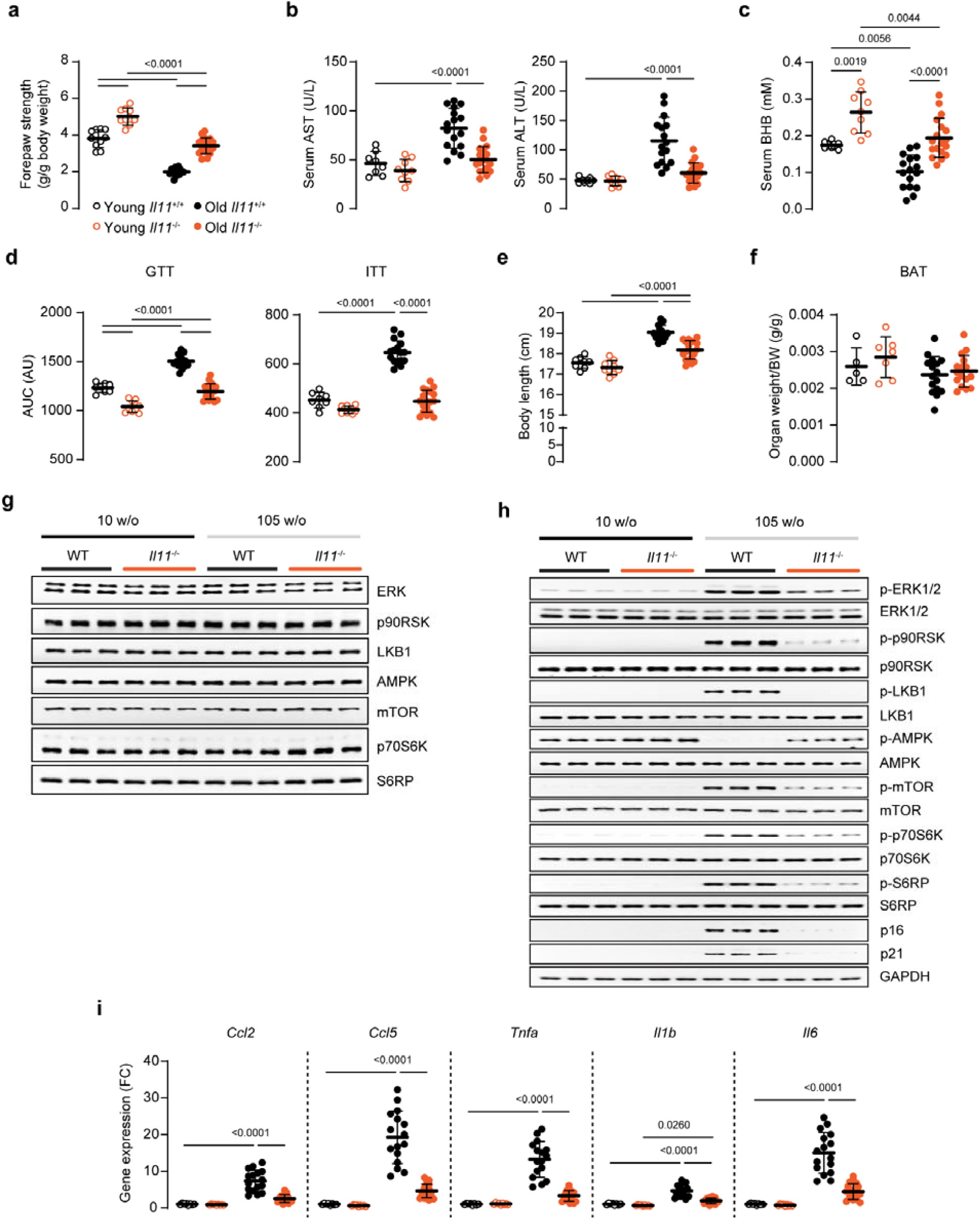
Female *Il11^-/-^* mice are protected from age-associated frailty and inflammation and have better metabolic profiles. **a** Front paw grip strength, serum levels of **b** ALT, AST, **c** BHB, **d** area under the curves (AUC) of glucose tolerance tests (GTT) and insulin tolerance tests (ITT), **e** body lengths, **f** indexed brown adipose tissues (BAT) weight, **g** WB of total proteins for the respective phospho proteins shown in **Fig. 2p** (representative datasets from n=6/group), **h** WB showing ERK1/2, mTOR, p70S6K, and S6RP activation and p16, p21, and GAPDH protein expression levels (representative datasets from n=6/group), **i** relative pro-inflammatory gene expression (*Ccl2, Ccl5, Tnf*lZ*, Il1*□ and *Il6*) levels in vWAT from young and old female WT and *Il11^-/-^* mice. **a-f, i** Data are shown as meanL±LSD, two-way ANOVA with Sidak’s correction. **a-e, i** young WT, n=8; young *Il11^-/-^*, n=9; old WT, n=16; old *Il11^-/-^*, n=18; **f** young WT, n=5; young *Il11^-/-^*, n=7; old WT and *Il11^-/-^*, n=16/group. FC: fold change; AU: arbitrary units.

**Supplementary Fig 5.**
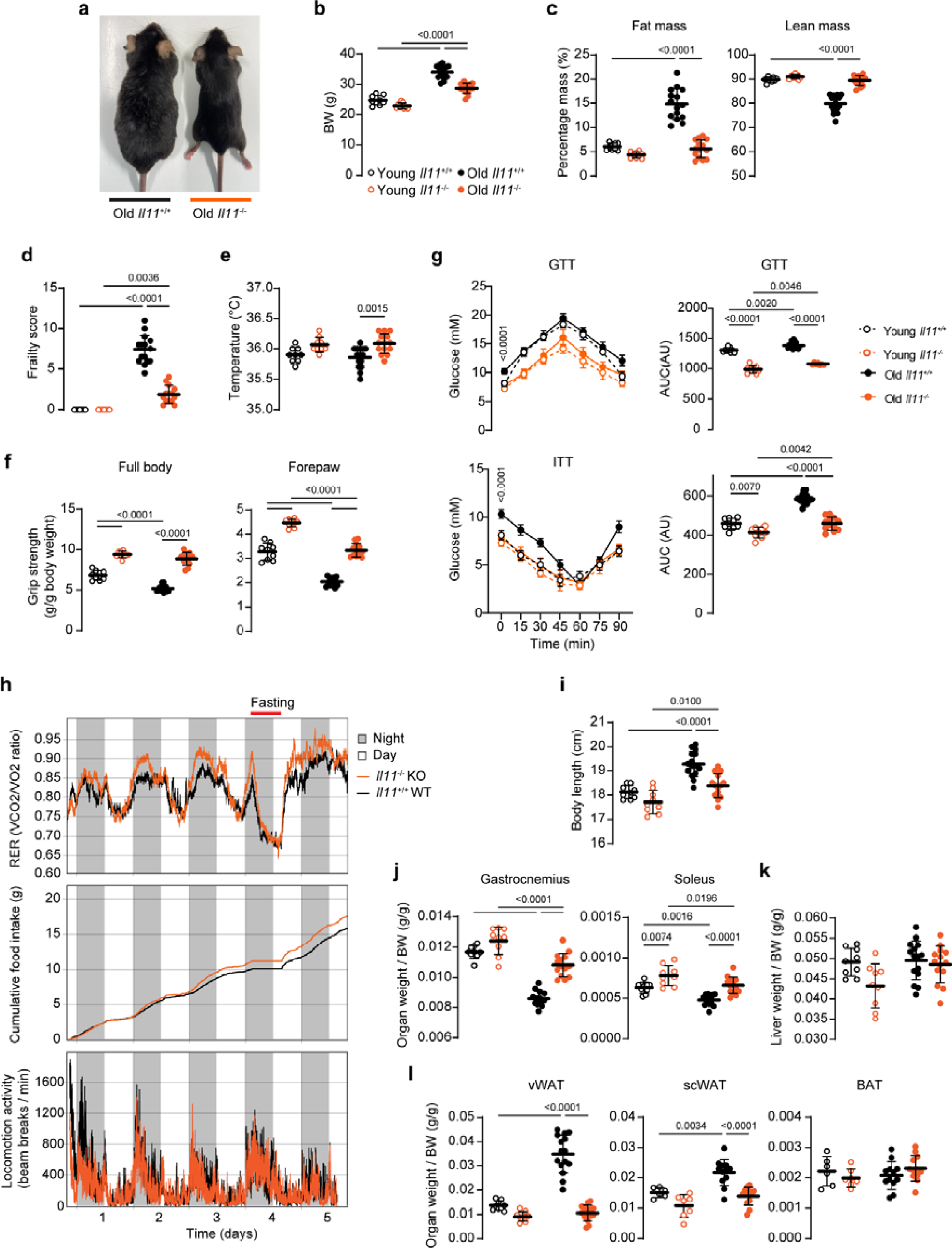
Old male *Il11*^-/-^mice are protected from metabolic decline. **a** Representative image of 108 w/o mice, **b** body weights (BW), **c** percentages of fat and lean mass (normalised to BW), **d** frailty scores, **e** body temperatures, **f** full body and forepaw grip strength measurements, **g** glucose and insulin tolerance tests (GTT and ITT) from young (12 w/o) and old (105 w/o) male WT and *Il11^-/-^* mice. **h** RER measurements, cumulative food intake, and locomotive activities as measured by phenomaster for 5 days (n=10/group) in 68-70 w/o male WT and *Il11^-/-^* mice (n=10/group). **i** Body length, **j** indexed weight of skeletal muscle (gastrocnemius and soleus), **k** liver, **l** vWAT, subcutaneous WAT (scWAT), and BAT. **b-g, i-l** Data are shown as mean☐±☐SD (young WT and *Il11^-/-^*, n=6-9/group; old WT, n=15; old *Il11^-/-^*, n=12-14), two-way ANOVA with Sidak’s correction. FC: fold change; AU: arbitrary units.

**Supplementary Fig 6.**
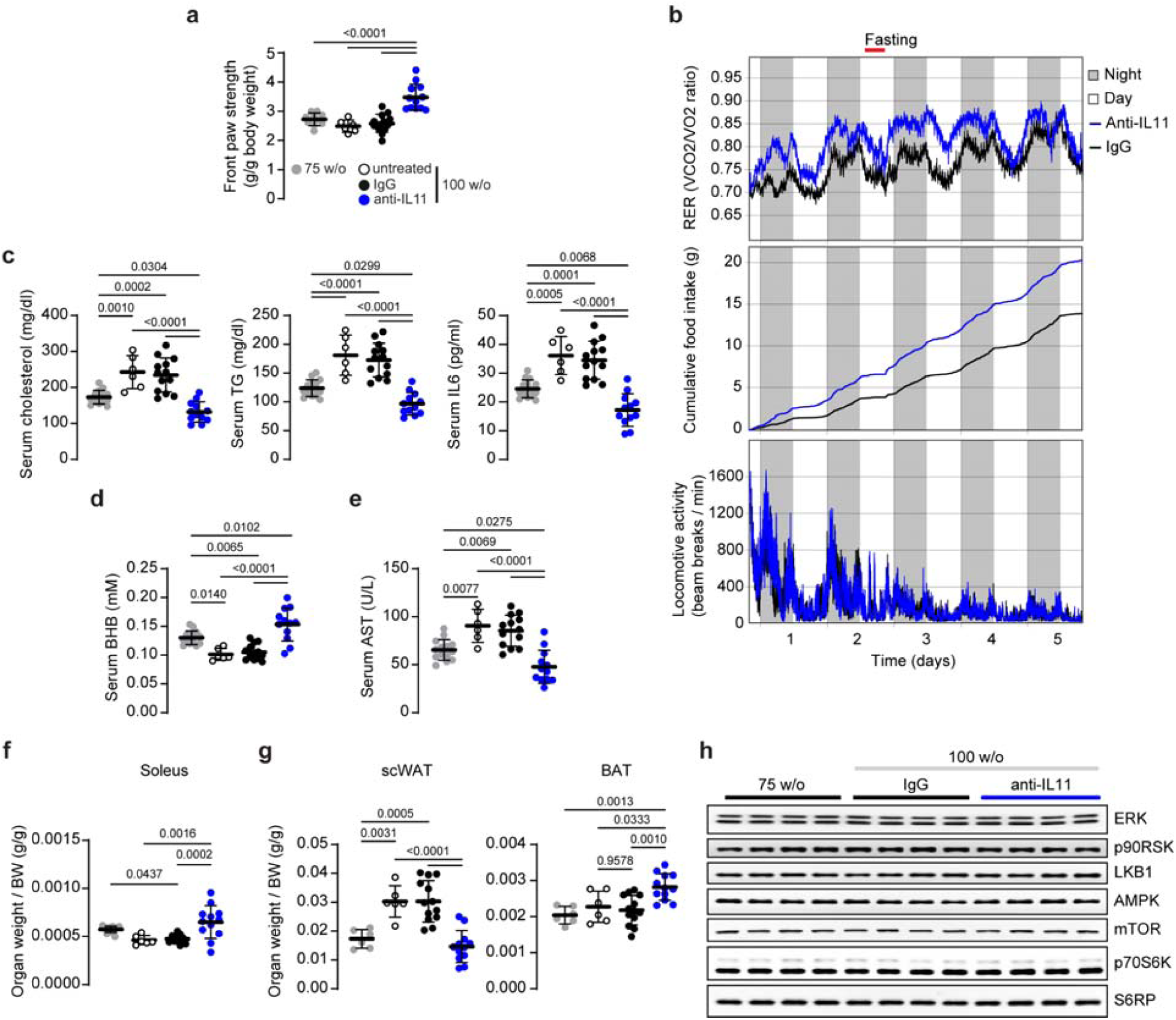
Anti-IL11 improves muscle strength and metabolic health. **a** Front paw grip strength, **b** RER measurements, cumulative food intake, and locomotive activities as measured by phenomaster for 5 days on IgG/X203-treated old (81 w/o) mice - 6 weeks after IgG/X203 administration was started (n=10/group), serum levels of **c** cholesterol, TG, and IL6, **d** BHB, and **e** AST, indexed weight of **f** soleus, **g** subcutaneous white adipose tissues (scWAT) and BAT, and **h** WB of total proteins for the respective phospho proteins shown in **Fig. 3m** (representative datasets for n=6/group) for anti-IL11 therapeutic dosing experiment in old mice as shown in Schematic **Fig. 3a**. **a, c-g** (75 w/o control, n=6-14; untreated 100 w/o, n=6; IgG-treated 100 w/o n=13; X203-treated 100 w/o, n=12). **a, c-g** Data are shown as meanL±LSD; one-way ANOVA with Tukey’s correction.

**Supplementary Fig 7.**
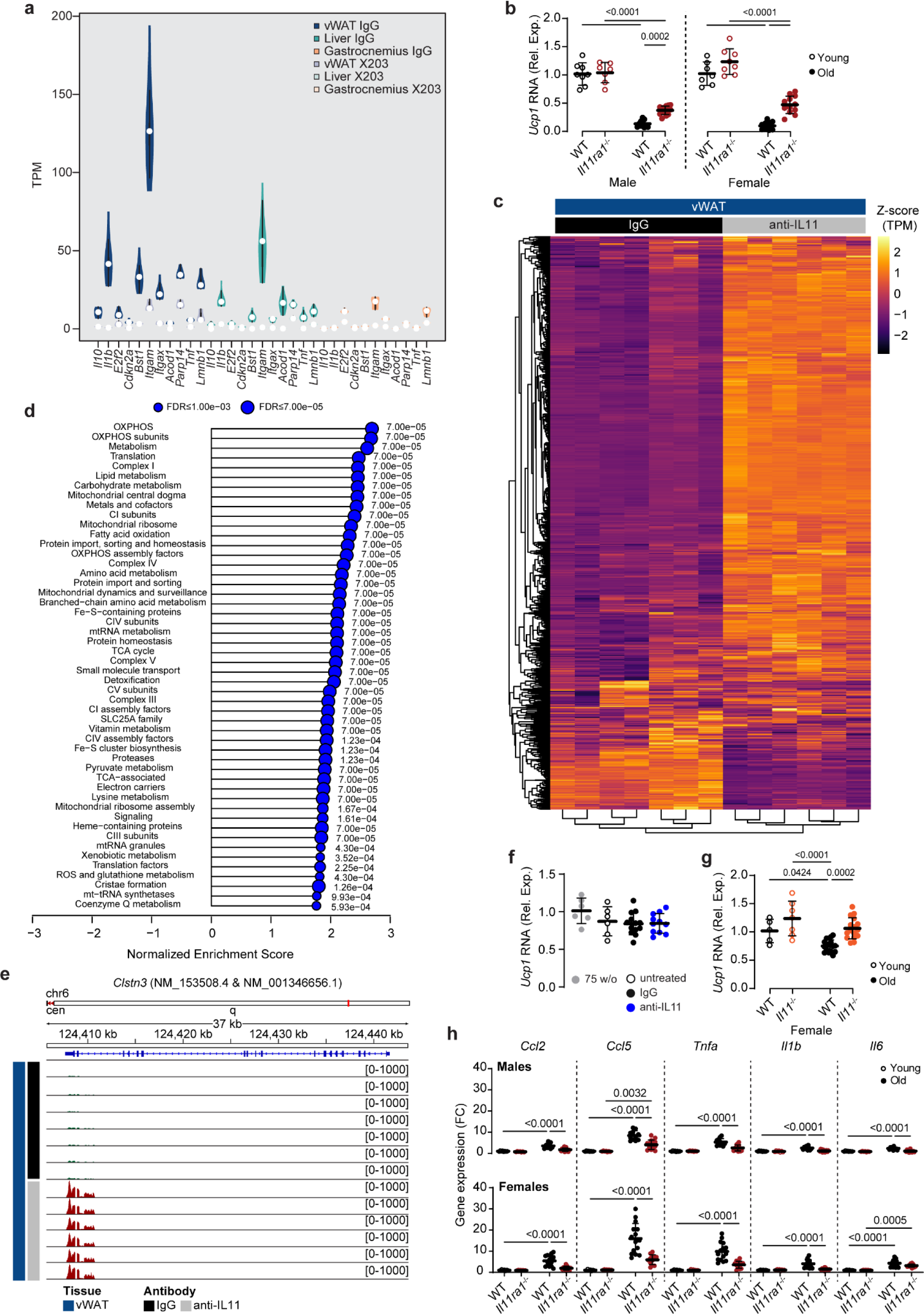
Beneficial effects of anti-IL11 in aged vWAT. **a.** Violin plot of Transcripts per million (TPM) values of senescence genes (based on Tabula Muris Senis consortium) in vWAT, liver, gastrocnemius samples from mice receiving either IgG or anti-IL11 as shown in schematic **Fig. 3a**. **b.** Relative *Ucp1* mRNA from young (10 w/o) and old (110 w/o) male and female WT and *Il11ra1^-/-^* mice. **c.** Heatmap showing row-wise scaled TPM values for the gene-list in Mitocarta 3.0. (no. of genes = 1,019 with TPM>=5 in at least one condition). **d.** A lollipop plot for top 50 Mitocarta 3.0 pathways found significant (p-adj < 0.05) in enrichment analysis using fgsea R package. No negative NES was found to be significant. **e.** Distribution of RNA-seq reads at the *Clstn3* locus from IgG or anti-IL11-treated vWAT. Relative *Ucp1* mRNA expression levels in BAT from **f** therapeutic anti-IL11 dosing group (left; 75 w/o control, n=6; untreated 100 w/o, n=6; IgG-treated 100 w/o n=13; X203-treated 100 w/o, n=12) and from **g** WT and *Il11*^-/-^ mice (right; young WT, n=5; young *Il11^-/-^*, n=7; old WT and *Il11^-/-^*, n=16/group). **H** Relative vWAT mRNA expression of pro-inflammatory markers (*Ccl2*, *Ccl5*, Tnf⍺, Il1flJ, *Il6*) in young (10 w/o) and old (110 w/o) male and female WT and *Il11ra1^-/-^* mice. **a, c, d, e** Liver and gastrocnemius (n=8/group), vWAT IgG, n=7; vWAT anti-IL11, n=6; **b, h** young male WT, n=8; young male *Il11ra1^-/-^*, n=7; old male WT, n=11; old male *Il11ra1^-/-^*, n=13-14; young female WT, n=7; young female *Il11ra1^-/-^*, n=8; old female WT, n=15; old female *Il11ra1^-/-^*, n=12. **a** Data are shown as violin plots with medianL±Lmin-max; **b, f-h** data are shown as meanL±LSD. **b, g, h** Two-way ANOVA with Sidak’s correction; **e** one-way ANOVA with Tukey’s correction.

**Supplementary Fig 8.**
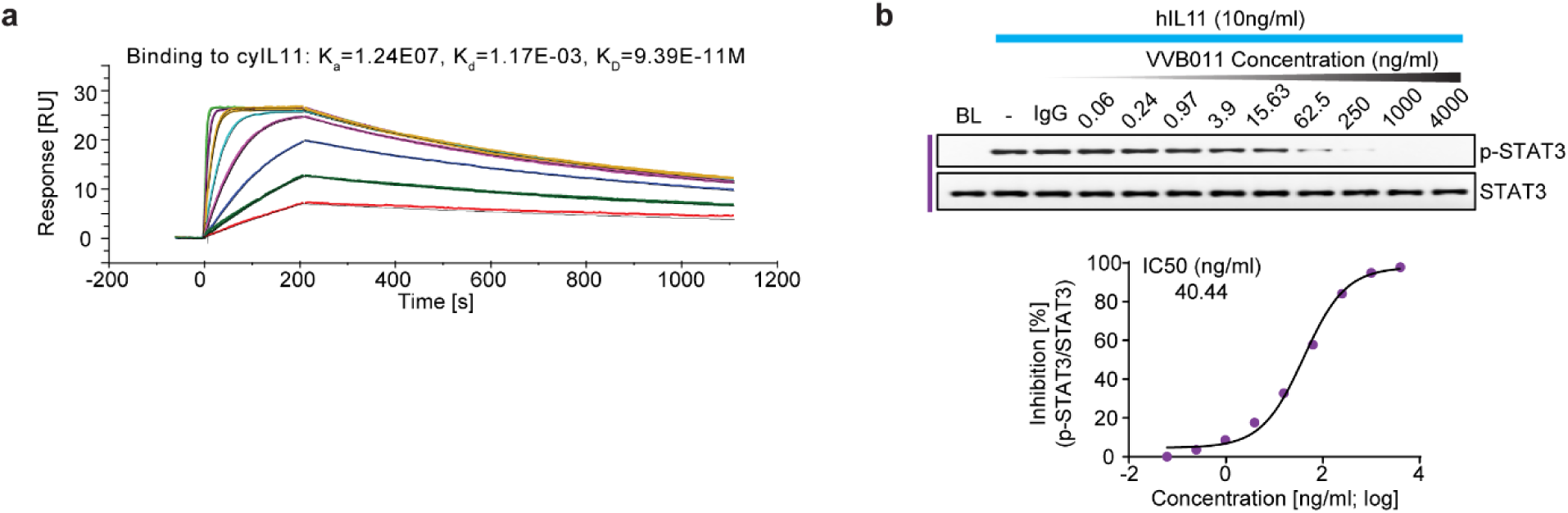
Development of a humanised, neutralising IL11 antibody. **a** VVB011 interaction with cynomolgus IL11 as determined by surface plasmon resonance (SPR). **b** Dose-response curve and half maximal inhibitory concentration value of VVB011 (61 pg/mL to 4 μg/mL; 4-fold dilution) in inhibiting STAT3 activation (by WB) by hIL11-stimulated A549.

**Supplementary Table 1.**
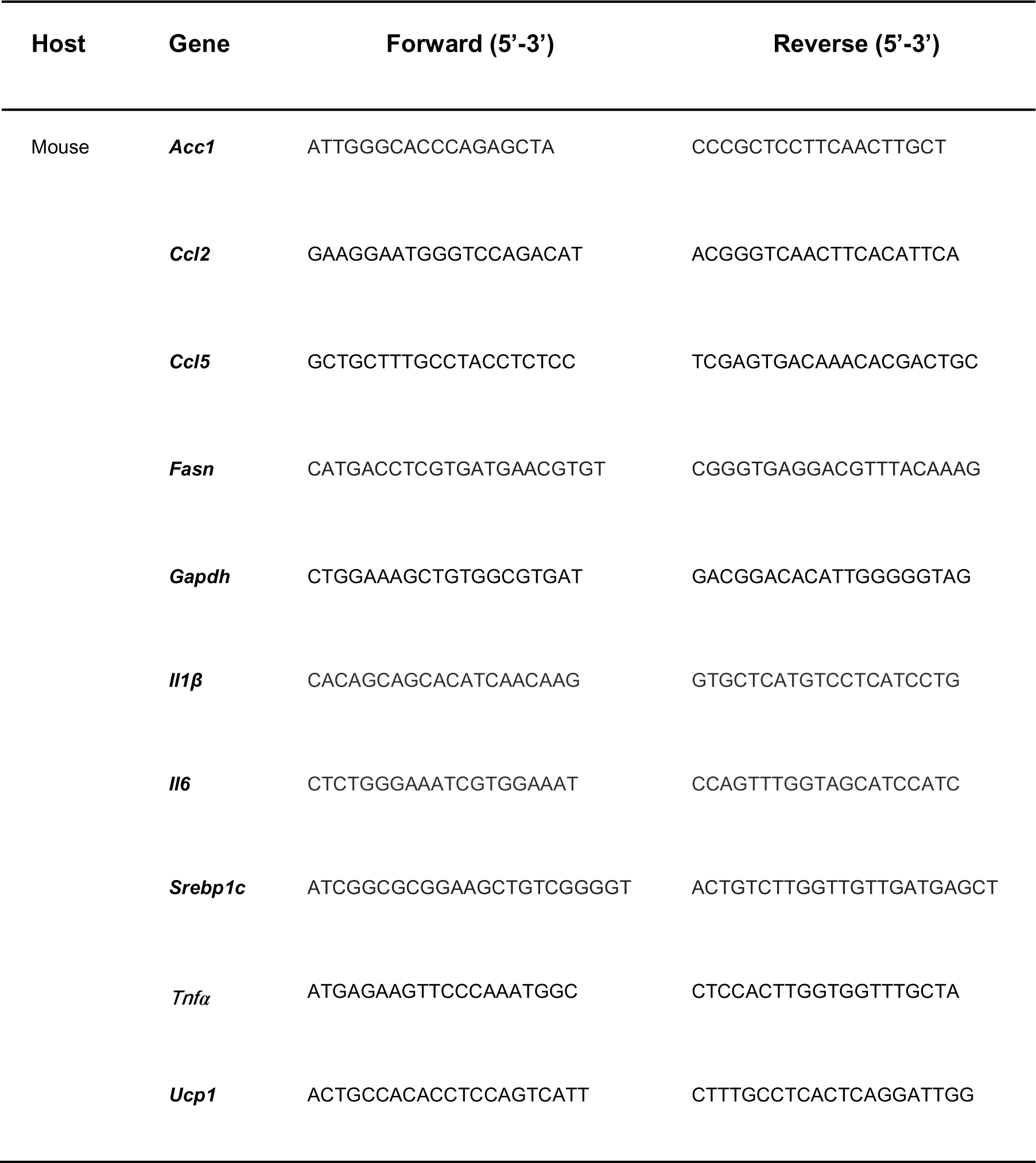
**List of primer sequences**

